# Reduction of SEM charging artefacts in native cryogenic biological samples

**DOI:** 10.1101/2024.08.23.609373

**Authors:** Abner Velazco, Thomas Glen, Sven Klumpe, Avery Pennington, Jianguo Zhang, Jake LR Smith, Calina Glynn, William Bowles, Maryna Kobylynska, Roland A. Fleck, James H. Naismith, Judy S Kim, Michele C. Darrow, Michael Grange, Angus I Kirkland, Maud Dumoux

## Abstract

Scanning electron microscopy (SEM) of frozen-hydrated biological samples allows imaging of subcellular structures at the mesoscale in their native state. Combined with focused ion beam milling (FIB), serial FIB/SEM can be used to build a 3-dimensional picture of cells and tissues. The correlation of specific regions of interest with cryo-electron microscopy (cryoEM) can additionally enable subsequent high-resolution analysis. However, the adoption of serial FIB/SEM imaging-based methods is limited due to artefacts arising from insulating areas of cryogenically preserved samples. Here, we demonstrate the use of interleaved scanning to reduce charging artefacts, allowing the observation of biological features that otherwise would be masked or perturbed. We apply our method to samples where inherent features are not visible. These examples include membrane contact sites within mammalian cells, visualisation of the degradation compartment in the algae E.gracilis and observation of a network of membranes within different types of axons in an adult mouse cortex. We further propose an alternative scanning method that could also be widely applicable to imaging any non-conductive.

## Introduction

Volume electron microscopy (EM) comprises a group of electron imaging techniques that can be used to visualise samples at different resolutions. The spectrum of techniques generally incorporates imaging using either a transmission electron microscope (TEM) or a scanning electron microscope (SEM). For TEM imaging, the main restrictions are related to the sample thickness, which is limited by the electron mean free path ^1,2^. The use of SEM imaging gives lower resolution but can be used to image larger volumes when combined with sectioning techniques using either a diamond knife or a focused ion beam^3^. However, with SEM imaging, samples can be subject to electrical charging while scanning, leading to the presence of undesirable artefacts in the images^4–6^. This charging depends on the sample preparation since the electrical conductivity can be modified by embedding or staining the sample^7,8^. Gas injection-based charge compensation and variable pressure SEMs have been used to reduce charging^9–11^ but have a negative impact on the spatial resolution and are incompatible with experiments under cryogenic conditions.

SEM acquisition parameters directly impact the dynamic charge-discharge behaviour in insulators^12^, and can be optimised to reduce charging effects for a particular sample^13^. For example, reducing the accelerating voltage (to find a crossover energy at which the deposited and emitted charge are balanced) has been shown to reduce the buildup of charge ^14,15^. Alternatively, reducing the dwell time (irradiation time per pixel) and increasing the distance between scanning points can also mitigate charging artefacts^16–18^ by reducing the local rate at which the charge is deposited in the sample. However, all these approaches have limitations. Reducing the voltage, beam current or the dwell time can reduce the signal to noise ratio (SNR) in the images and may require the use of dedicated electron optical and detection systems, whilst increasing the distance between scanning points by simply reducing the magnification lowers the resolution.

In many cases, images are not acquired with the full electron fluence (electrons per unit area) per scanning position in one pass. Instead, the fluence is distributed across several line scans or in multiple frames before integration of the signal^16,18^. Integration of the signal line-by-line avoids potential issues with drift but deposits the electron fluence in a limited area (a line), which can lead to charge build up and cause directional charging artefacts^19^. Frame integration is also subject to drift due to the increased time required to scan the same position and also requires alignment of successive frames.

For insulating samples, these steps are often not sufficient to avoid charging artefacts^19–21^. Hence efforts have been made to computationally correct for charging artefacts, by using predictive models to subtract these and to recover the underlying signal^19,22^. However, directly reducing charging artefacts during imaging is preferable. Importantly, biological samples are inhomogeneous, adding further complexity to optimising beam and imaging conditions. The most common charging artefacts observed in vitreous biological samples are dark streaks around high-contrast structures such as lipid droplets (LD) or myelinated axons^19,23^, and inhomogeneous contrast manifested as extremely dark and bright pixels^20^. Importantly, existing schemes that reduce charging artefacts in one region of an inhomogeneous sample may reduce the SNR or sharpness in other regions that do not experience these effects.

Here we describe a new approach to mitigate charging artefacts by using an interleaved, or ‘leapfrog’, scanning pattern^24^. This approach distributes the electron fluence differently, in time and space, compared to conventional raster scanning, and allows for a more efficient charge dissipation. By applying image registration correction (MotionCor2^25^) to correct for beam and mechanical induced motion, frames from sequential acquisitions can be aligned and integrated to improve the SNR. Using different biological samples, we compare image acquisition strategies that demonstrate improvements in image quality; raster scans with frame integration, raster line integration which is a common scanning strategy in SEM imaging^19,23,26–28^ and interleaved scans with frame integration. We demonstrate that interleaved scanning allows the imaging of cells and tissue in a near-native state with minimal charging artefacts. The use of interleaved scanning also enables the observation of biological features often obscured by charging artefacts, specifically diverse membrane contact sites (MCS) in a mammalian cell model, the complex layers of the axon in mouse brain and the organisation of the network of degradative compartment in *E.gracilis*.

## Results

### Experimental set up

SEM images are typically acquired using raster scan patterns, where a line is imaged in one direction, often the x axis of a rectangular coordinate system, typically from left to right (Figure 1A and C), followed by the acquisition of the next line with a shift along the y axis. In this array, the interval between each x position is equal to the dwell time (fast scanning direction) while in y (slow scanning direction), the interval is equal to the product of the dwell time and number of pixels along the x axis. In contrast, interleaved scanning skips adjacent pixels in the x and y scan directions. Specifically, the interleaved scan pattern used in this work skips 2 pixels in the x direction, returns to the starting x coordinate while skipping 2 pixels in the y direction and repeats until all positions in the frame have been visited (Figure 1B and D). This interleaved configuration allows a longer time interval between subsequent x or y positions than for an equivalent raster scan.

**Figure 1:**
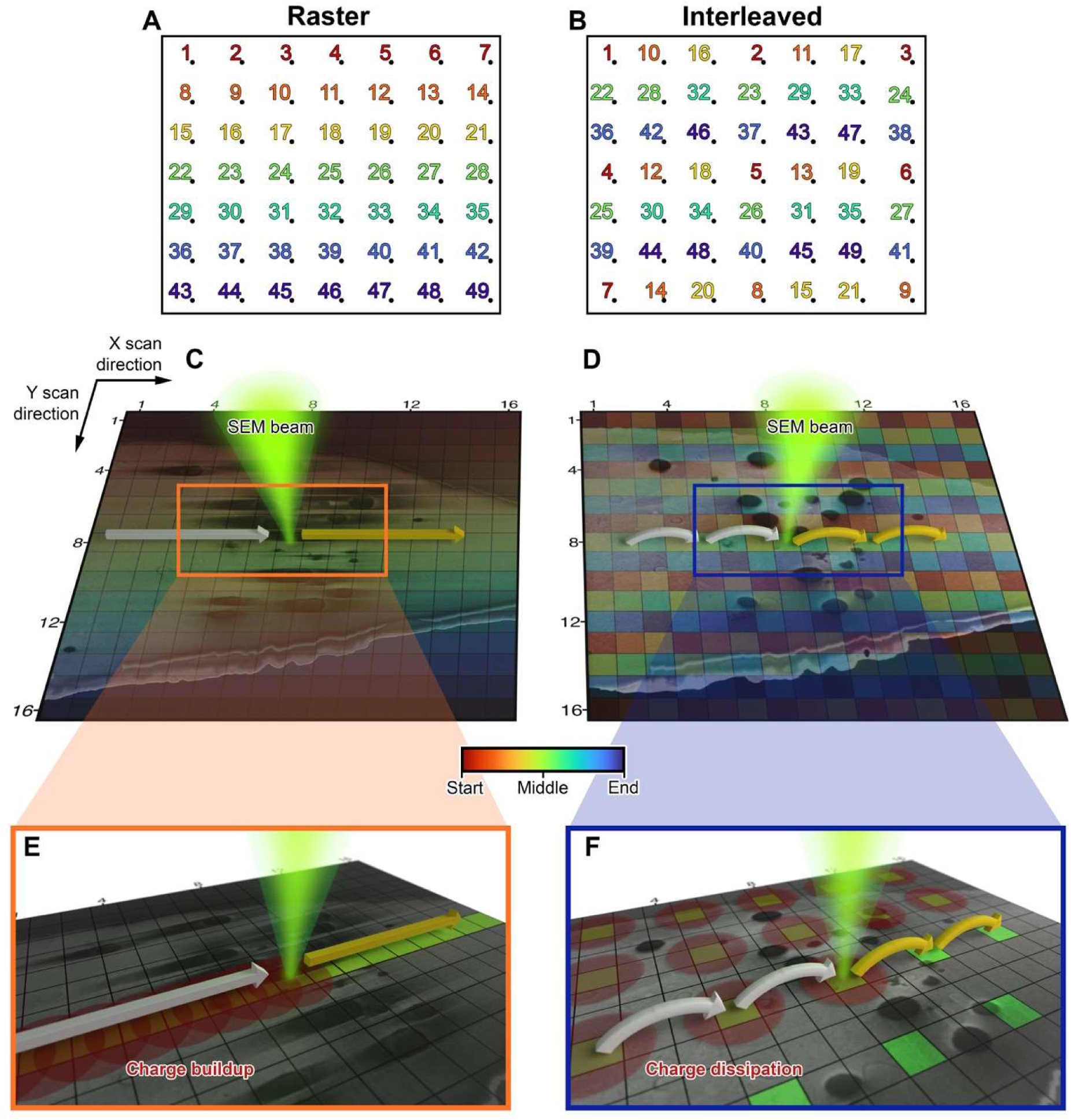
Schematic showing different scanning patterns. (A and B) a field of view (black rectangle) imaged using a 7×7 array where the rainbow colour corresponds to the temporal distribution of the fluence with red at the start and blue at the end of the time sequence. (A) Raster scans visit each x position before jumping to the next y position. (B) Interleaved scans which skip 2 pixels in x, before and 2 positions in y. (C-F) Overlay of a biological sample (here vitrified RPE-1) with a schematic representation of pixel positions (black squares) in a 16×16 array. The green cone represents the electron beam. The straight arrows represent a raster scan while the curved arrows indicate pixel skipping during interleaved scanning. The colour of the arrow represents the pixel acquired (white) or to be acquired (yellow). (E and F) spatial distribution of the fluence for isotropic charge dissipation. Yellow pixels have been imaged, green pixels are to be imaged. Red circles illustrate charge dissipation radiating outwards at each position.

The spatio-temporal fluence distribution (raster or interleaved) is convolved with the averaging mechanism used to form an image. We hypothesise that in 2 dimensions the spatial charge distribution is bigger than the beam size (estimated to be a 2 nm FWHM in our experiments) and that the charge dissipates uniformly in all directions from this point, assuming the sample has isotropic electrical properties^29–31^ (Figure 1E and F). In this model, line integration accumulates charge locally as the beam is constantly rastered along a line, and hence charge can dissipate more efficiently in the y direction but not in the x line direction as this is constantly scanned (Figure 1E). For frame integration, the charge does not accumulate in a local area but accumulates across the entire frame as charge dissipating from one position will accumulate during the exposure of the neighbouring position. Hence, for a raster scan the charge dissipation is not homogeneous whereas for interleaved scanning, the charge dissipation is more uniform along the x and y axes (Figure 1F), which is consistent with a previous report that shows a reduction in effects due to charging and beam damage^32^ for room temperature SEM and scanning-TEM^17,33^. The interleaved scan described in this work demonstrates one of many options to optimally engineer data collection strategies for cryogenic SEM specimens.

As biological exemplars, we report data from a mammalian cell model (RPE-1), a single cell algae (*E.gracilis*) and mouse cortex. The mammalian cells and algae were vitrified by plunge freezing and the tissue was vitrified by high pressure freezing (HPF). No samples were chemically fixed, dehydrated or stained. For plunge freezing, contact with a thermally and electrically conductive support (gold or carbon) is important for charge dissipation during imaging^34,35^. In contrast, during HPF of tissue, a thick tissue biopsy was deposited in a gold coated copper carrier filled with cryoprotectant before vitrification. Thus, the sample has limited direct contact with the carrier, reducing the electrical conductivity and the dissipation efficiency during imaging.

Diverse biological samples of inherently different molecular assemblies as well as varied sample preparation methods are explored to test if engineered scans are beneficial. Previous reports show that some components in the cell are more prone to charging artefacts^19,23,26,27^. Lipid droplets, stacking of membrane (thylakoid membrane in chloroplast, myelin sheets in brain) and degradation compartments (endosomes and lysosome) are often local charging centres due to their molecular composition. Using different scanning strategies, we confirm that charge redistribution and longer dissipation times can mitigate SEM charging artefacts.

### Scanning pattern and dwell time

In our previous work^19^ we demonstrated that using an Ar plasma for sputtering while imaging at low kV (∼1 kV) can be used to create volumes of vitrified biological samples at ∼10nm resolution. Based on this work, we set a constant electron fluence (10^−2^ e^-^/Ǻ^2^) using three combinations of dwell time and number of scan repetitions to vary the pixel flux in three regimes defined as: low (100 ns x100 repetitions), intermediate (500 ns x20 repetitions) and high (1000 ns x10 repetitions). The repetitions were completed either at the line integration (LI) or frame integration (FI) level and the scan pattern was either raster or interleaved. The angle between the electron beam and the sample dictates the image quality. When imaging at an angle, often 52°, the total secondary electron yield is optimal for the instrument used but differences in the electrical properties of the sample are lost leading to a reduction in the image contrast ^36^. Consequently, to optimise the image contrast, by analogy with our previous work^19^, imaging perpendicular to the milled surface increases the contrast^36^. As both of these imaging geometries are commonly used, we imaged the samples with the SEM column at 52⁰ (Figure 2 and Extended Data Figures 1-3) and 90⁰ ( Extended Data Figures 5-7) relative to the milled surface of the sample. When the region of interest (ROI) was exposed to 10^-2^ e^-^/Å^2^, the surface was polished with an Ar plasma with a 50 nm step to avoid accumulation of electron beam damage during multiple acquisitions.

**Figure 2:**
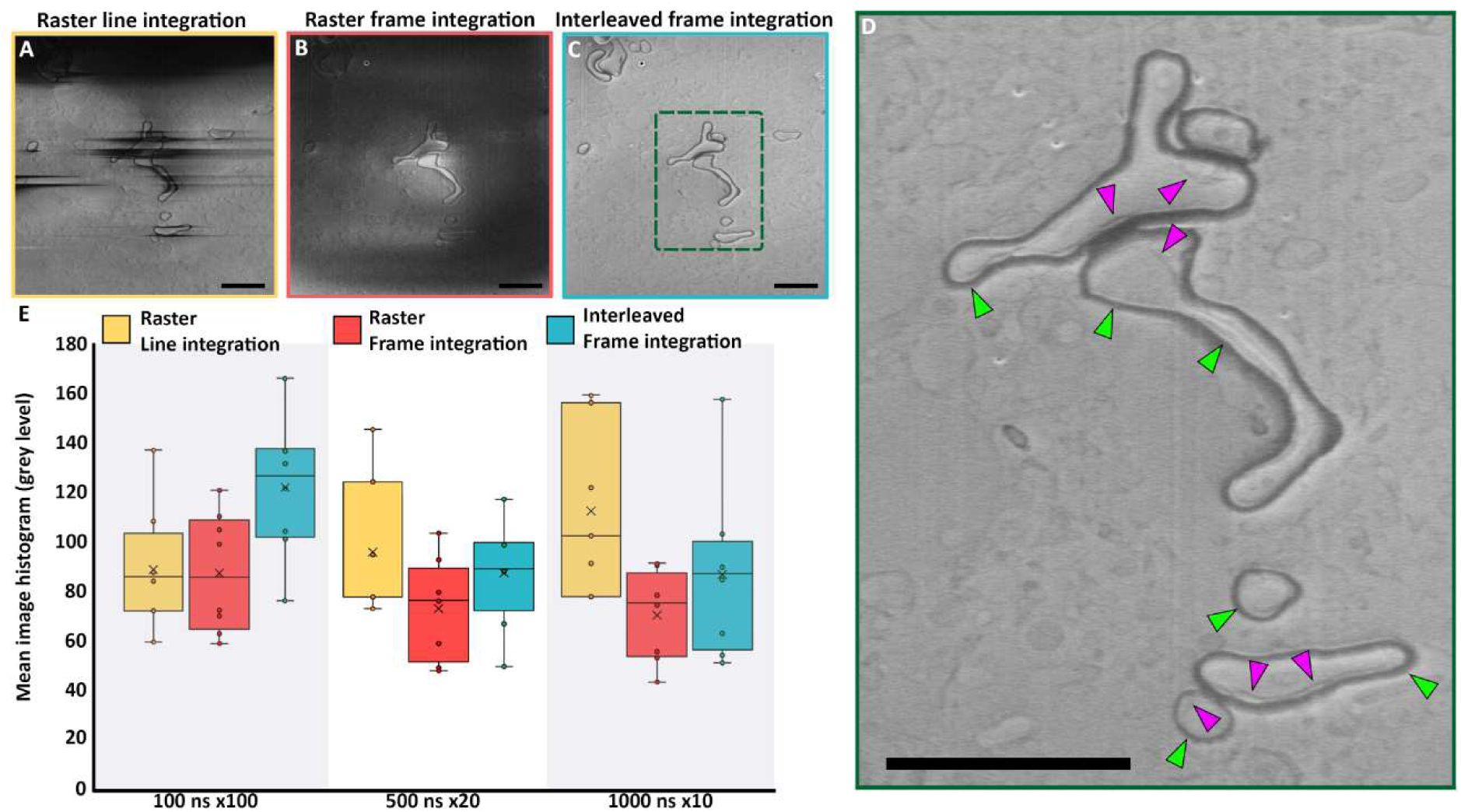
Effect of electron fluence distribution on charging artefacts (SEM imaging at 52° angle to the FIB milled sample plane). **(A-C)** Representative images of brain tissue for different pixel fluence distribution strategies with raster or interleaved scan patterns **(D)** Enlarged area highlighted in **C** by a green dashed rectangle that shows detail visible within the membranous layers of myelination in the brain. Green arrows indicate the myelin sheath while pink arrows indicate the oligodendrocyte inner tong. **(C and D)** Interleaved frame integration at 100 ns × 100 shows the greatest improvement in reducing charging artifacts. Scale bar: 2 µm. **(E)**: Mean of image intensity histograms of different images (population n>5) from vitrified RPE-1, *E.gracilis* and mouse brain images represented as circles. Population median (middle line), population mean (cross), median of the bottom half (bottom box), median of the top half (top box). Vertical lines extend to minimum and maximum intensity values. Statistical analysis is given in Supplementary Table 1. Images acquired using interleaved scanning with frame integration using a short dwell time and high integration (100 ns × 100) form a dataset for which the mean intensity is 122, closest to 127, which is the mid-range histogram intensity value of the 8-bit image data (0-255 range). Deviation in intensity from this value indicates charging as described in the text.

Our data show that for all fluence distributions and imaging angles considered, images acquired using raster scanning with line integration are dominated by streaks in the fast scan direction (Figure 2A. Extended Data Figures 1A-C, 2A-C, 3A-C,, 5A-C, 6A-C and 7A-C). However, an increase in dwell time (1000 ns) improved only some patches within the image where biological features could be discerned without artefacts (Extended Data Figures 1C, 2C and 3C), highlighting the difficulty that can arise when optimising SEM imaging conditions for inhomogeneous samples. Using frame integration with a raster scan pattern, images obtained at a glancing angle are corrupted by charging artefacts in the form of dark areas covering the majority of the scanned ROI. For longer dwell times (500 ns and 1000 ns), localized bright charging artefacts were also observed (arrows in Extended Data Figures 1E-F, 2E-F, 3E-F and 5E-F, 6E-F). When using an interleaved scan pattern in combination with frame integration we observed that the charge accumulation is dependent on the dwell time irrespective of the imaging angle, with a 100 ns dwell time image showing minimal charging artefacts (Figure 1C and Extended Data Figures 1G-I, 2G-I, 3G-I, 5G-I, 6G-I and 7 G-I). In comparison to raster scanning, HPF vitrified mouse cortex imaged with an interleaved scan shows multiple membrane compartments isotropically resolved within regions that could previously not be seen. This included the whole myelin sheath (green arrow figure 2 D) and internal membrane compartments (the inner tong) (pink arrow Figure 2 D). Being able to resolve the internal membrane near the lipid-rich myelin which is prone to charging demonstrates the improvement from interleaved scanning for reducing charge artefacts enabling the visualisation of additional details within frozen hydrated samples. Other imaging strategies (Figure 2A-B) clearly result in significant artefacts that obscure these details. We further note that previous work investigating the myelin sheath have required resin embedding and staining^37–40^ whereas we are able to image natively preserved frozen-hydrated samples.

The quality of an image can be characterised from the pixel intensity histogram (Extended Data Figure 4). Typically, an image with minimal charge artefacts and high contrast features of interest would have a mid-range Gaussian distribution of intensity 127 for 8-bit images (Extended Data Figure 4G as an example). However, any accumulation of charge as extreme dark or extreme bright pixels causes an asymmetry in the distribution (Extended Data Figure 4 A-F and H-I). We have compiled our data and extracted for each image the mean of the histogram. Some variation in the mean from sample to sample is expected but if the whole dataset consists of high quality images, the population concentrates around the mid-range scale. Deviation from this implies that the dataset is affected by large numbers of extreme pixel intensities, resulting from poor image quality. Irrespective of the imaging angle, the use of interleaved pattern with frame integration using the short dwell time fluence deposition strategy performed best with a population centring around a 122 grey level when imaging 52° to the milled surface (Figure 2E) and 97 when imaging perpendicular to milled surface (Extended Data Figure 5). All the other imaging conditions give datasets with a lower mean (Figure 2E and Extended Data Figure 5J). We hence conclude that imaging with interleaved scanning using a short dwell time reduces charging artefacts, irrespective of sample surface angle and the biological sample types examined.

### Scanning strategy for automated image segmentation

Biological volume data acquired using SEM images is often post-processed using segmentation algorithms to extract insights relevant to a specific biological question (for example tissular/cellular organisation, volume analysis or length of contact)^40–42^. However, curtaining^43,44^ and charging artefacts are often a source of poor segmentation outcomes^19,45^. Consequently, different approaches have been used to computationally remove both types of artefacts^46,47^. Hardware implementations, such as specialised sample stages (rock milling stage) and the use of a plasma FIB argon source, have also been employed to reduce curtaining artefacts^19,28,48–50^. We have investigated whether the implementation of interleaved scanning could benefit automated segmentation. We used the Segment Anything Model (SAM)^51^ on our shortest dwell time (100 ns) datasets for raster line/frame integration and interleaved frame integration with minimum user input and the same set of parameters. The number of picked objects were counted as well as the complexity of their shape. Our overall aim in this analysis is to observe the relationship between the reduction of charging artefacts and unsupervised segmentation rather than to obtain the optimum segmentation for a defined component. This analysis was applied to mouse brain tissue, RPE-1 cells and *E.gracilis* imaged at 52⁰ or 90⁰ with respect to the milled sample plane. Extended Data Figure 8 shows a representative result obtained from the segmentation. The absence of artefacts in data acquired using interleaved frame integration scanning allowed more objects to be picked (Extended Figure 8A-F). These objects were not deformed or attributed to charging artefacts (as is the case in Extended Data Figure 8A, arrows).

Datasets of RPE-1, *E.gracilis* and mouse brain acquired at the same angle were merged to measure the number of objects picked using SAM and their complexity defined as a ratio between the squared perimeter and the area. As each dataset has a unique biological content, the value of the number of objects picked or complexity score was normalised to that calculated from the corresponding interleaved frame integration. Interleaved scanning images systematically allowed SAM to identify and locate more objects than the counterpart rastered images (Extended Data Figure 8G). Imaging using raster scan frame integration produces images with artefacts that reduce the object picking by half on average. Only imaging using raster line integration at 52° allowed object picking with a value close to the reference (interleaved frame integration) with 80% of objects picked. In addition to the decrease in the number of objects picked by SAM when charging artefacts are present, the segmentation of those objects are often deformed from their actual shape^19,23,26,27,41,42^. Our analysis demonstrates that the objects picked by SAM from images acquired with raster imaging (frame and line integration) have a lower complexity score (20 to 40% less depending on the conditions), than those picked from images acquired with interleaved scanning (Extended Data Figure 8H). Overall, we conclude that a combination of interleaved scanning with short dwell times and frame integration allows the imaging of vitrified biological samples with minimal charging artefacts, subsequently enabling accurate segmentation of heterogeneous biological features.

### Lipid droplet membrane contact site morphological interactions

When imaging mammalian cells, the presence of LD often leads to very significant charging artefacts, and obscures the proximal detail of their local environment. Furthermore, using classical fixed, stained and embedding procedures for imaging at room temperature leads to artefacts, raising ambiguity about the identification of features and the relevance of the data^52,53^. Therefore, imaging vitrified samples is currently the most appropriate technique to study LD biology, but, as observed in Figure 3 and Extended Data Figures 2,3, 5 and 6, LD act as strong charging centers, making an analysis of their environment and interactions with other organelles difficult^19,26^. Using traditional raster methods (Figure 3D-E), the neighbouring environment around the LD is not clear, the ER (white arrows) is not visible, the mitochondria are not well defined. Furthermore the inside of the degradation compartment (black arrows) is heavily clouded by charging artefacts. With interleaved scanning it is possible to observe mitochondria distinctly including cristaes, possible ER membranes and a degradation compartment within the proximity of a LD (Figure 3F). Additionally, the image acquired using interleaved scanning allows the internal structure in the degradation compartment to be clearly observed as an interconnected network of electron dense material (black arrows). These observations are not possible or are severely limited in images acquired using other scanning strategies.

**Figure 3:**
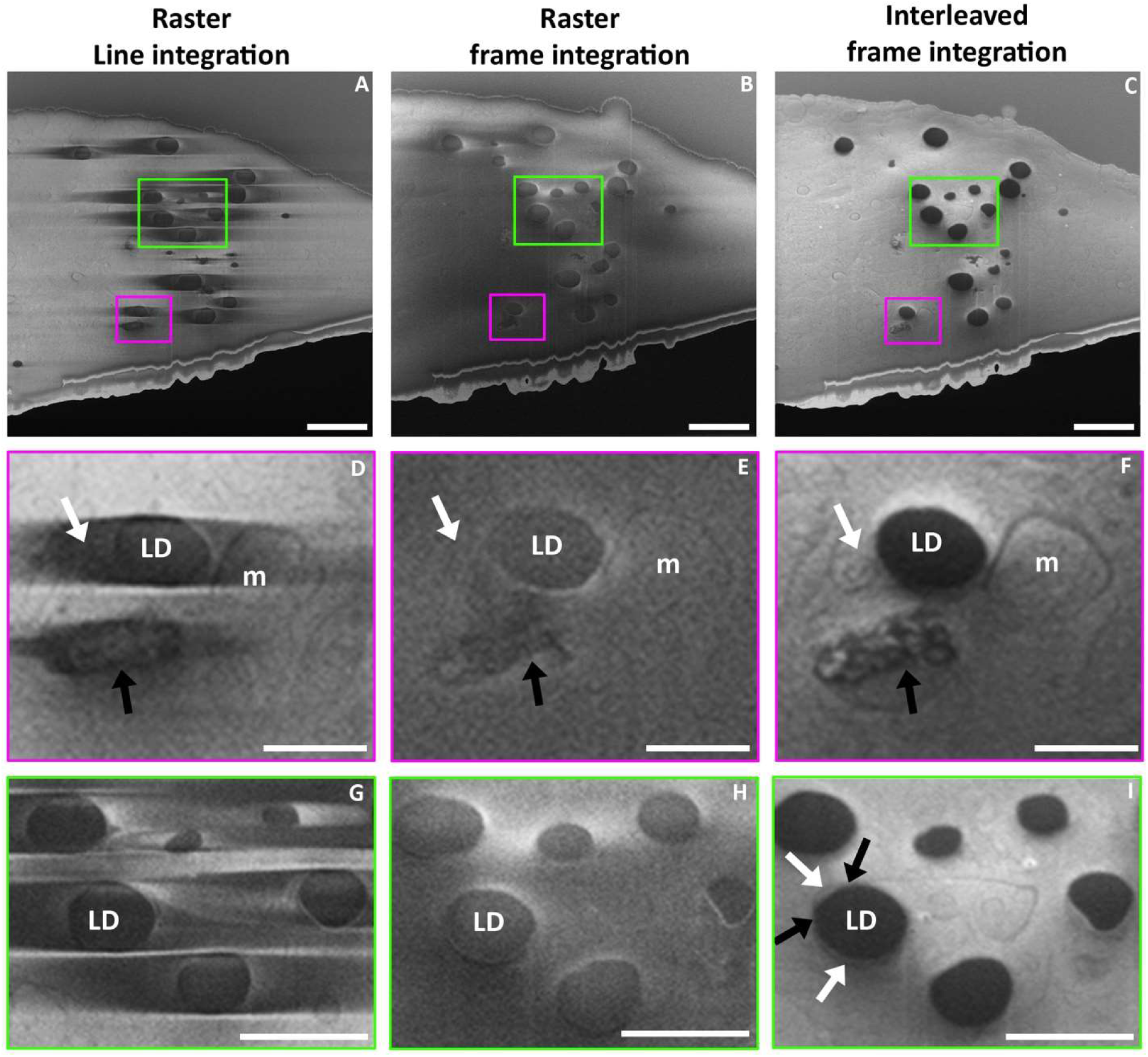
Charging artefact reduction in RPE-1 cells allows the observation of lipid droplet (LD) surrounding environments and LD to organelle membrane contact sites (MCS). Vitrified cells imaged at 52° with respect to the FIB milled sample plane using a 100 ns dwell time x100 repetitions. (A-C) Overview of slice of a RPE-1 imaged using raster line integration, raster frame integration and interleaved frame integration, respectively. Scale bar: 2 µm. (D-F) enlargement of the pink boxes in (A to C). White arrow: membrane in the vicinity of a LD. Black arrow: content of a degradative compartment. m: mitochondria. Scale bar 0.5 µm. (G-I) Enlargement of the green boxes in (A to C), brightness and contrast were modified to improve visualisation. White arrow: membrane contact sites (MCS) by juxtaposition of the endoplasmic reticulum (ER) and the LD membranes. Black arrow: membrane hemi fusion MCS between the ER and the LD. Scale bar 1 µm.

LD are critical lipid sources for organelles which form MCS with the surface of the LDs. The additional information in images formed using interleaved scanning allows the observation of subtle variations in the type of MCS (Figure 3G-I). As an example, two types of endoplasmic reticulum (ER)-LD MCS are observed (Figure 3I). The first, highlighted with white arrows, is characterised by a juxtaposition of membranes of the two organelles. Here, the membranes are tethered by protein-protein interactions and the transfer of material between the two membranes is mediated by a lipid transfer protein or a hydrophobic channel protein^54,55^. The other ER-LD MCS, highlighted with black arrows, is characterised by bridging contacts where the surface of the LD is continuous with the cytoplasmic leaflet of the ER allowing direct diffusion transfer of lipids between the two organelles^54,55^. In this example we have demonstrated that the alternative scanning strategy has a positive impact on studies of the interaction of LD with other organelles enabling the identification of subtle differences in MCS complexes in native conditions.

### Analysis of the degradation compartment in E.gracilis

Unicellular organisms, such as *E.gracilis,* can control complex functions, such as motility, hunting, and digestion through specialised sub-cellular structures. Hence observation of the relevant membranes of key sub-cellular compartments is critical to understanding how algae control these functions at a sub-cellular level. Extended Data Figure 9 shows a sectional plane near the anterior end of the algae, close to the insertion point of the flagella. Imaging using interleaved scanning improved the observation of features in the vicinity of the reservoir and surrounding structures (Extended Data Figure 9A-C), including paramylon and the membrane of the reservoir itself. We also observed that interleaved scanning is necessary to observe the thylakoid membrane sheets, the ER, neighbouring LD, vesicles contacting the reservoir as well as the membrane neighbouring the contractile vacuole (Extended Data Figure 9D). Even in areas where charging artefacts are less common (Extended Data Figure 9G-I, K-L), images recorded using interleaved scanning reveal additional details in the ultrastructure. For example, in Extended Data Figure 9J, the membranes separating the contractile vacuole and the reservoir are clearly visible and in Extended Data Figure 9K, the membrane delineating the degradation compartment is visible. Similarly, in Extended Data Figure 9L, the presence of LDs does not prevent the observation of the neighbouring chloroplast and associated thylakoid membrane.

It was further possible to reconstruct in three dimensions the network of degradation vesicles after volumetric acquisition (1600 µm^3^) of *E.gracilis* cells (Figure 4 and Supplementary movie 1). From observations of 55 vesicles only two are in proximity (<500 nm) to the reservoir and contractile vacuole, while the remainder form densely packed networks surrounded by paramylon and chloroplasts. Segmentation of the degradative compartment (DC), and its respective vesicular content, in one of the cells (Figure 4A) shows that all the DC are dissimilar. The DC is equivalent to the digestive system in multicellular organisms, and by understanding their structure, we can gain insight into their dynamic functionality. By plotting the volume of the vesicle as a function of the density of its content, (Figure 4B) we can distinguish the different vesicles populations (quadrant count ratio of 0.037). Most of these have a small volume (23/26 are under 1 µm^3^) and within this population, half have 20% or more of the total volume filled with dense content. This suggest that these different vesicles are at different states in their development. We have also observed the environment of these different populations and noticed the presence of extended contact between the nucleus, mitochondria or chloroplasts and the DC (Figure 4C and D). These contacts were mapped as it was possible to observe the deformation of the membrane of the organelles indicative of the presence of structured MCS between DCs and the chloroplast, nucleus or mitochondria. We did not observe specific extended contact with the Golgi apparatus, the reservoir or contractile vacuole. The DC forming these MCS are part of the low internal density population suggesting that the content of the DC may be used by other organelles. This hypothesis is supported by our observation of emptied vesicles forming MCS (Figure 4E) and we propose that these empty vesicles are emptied degradative compartments. Another specialised compartment that was characterised is the eyespot (Figure 4F-G). Eyespots are formed by carotenoid globules close to the reservoir, whose role is to detect and direct the cell depending on the light source, therefore allowing the appropriate amount of light for the algae to develop^56^. To date, studies of the eyespot have been limited to correlative approaches using fluorescence and resin embedded electron microscopy ^57^. Using volume imaging, we were able to image for the first time these structures and their integrity and model in 3-dimensions the organisation of their granules using a single approach (Figure 4F-G). Overall, in this aspect of the work reported we have demonstrated that by engineering the scanning strategy we could image single cellular organisms to analyse the intracellular organisation of key components of their life cycle.

**Figure 4:**
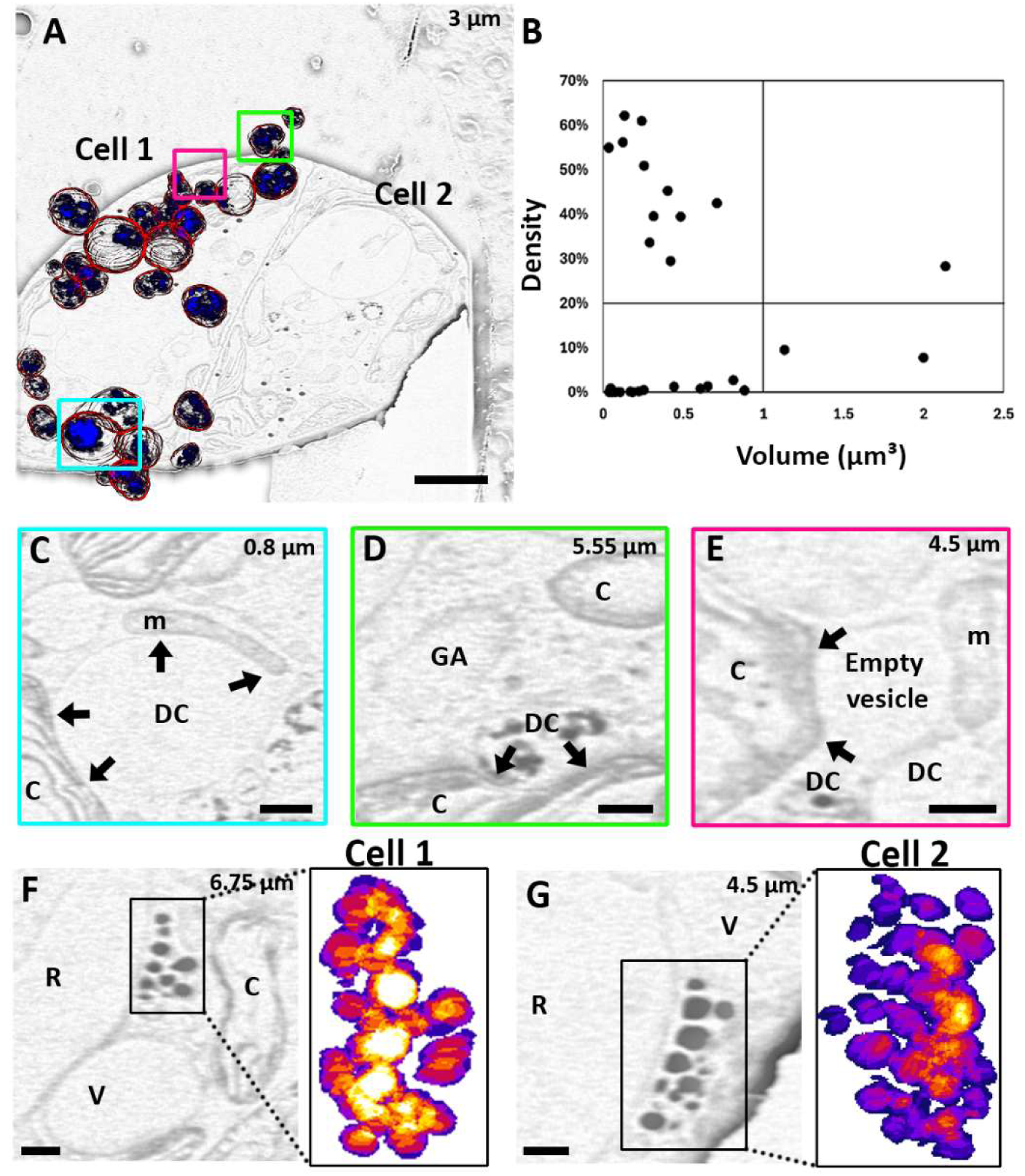
SEM volume imaging of *E.gracilis*. *E.gracilis* imaged at 52° with respect to the FIB milled sample plane using 100 ns dwell time x100 repetitions. A 1600 µm^3^ volume in focus aligned and subject to manual segmentation of the region of interest. (A, C-G) Images after background removal (see material and methods) to assist segmentation. For (A, C-G) the number in the upper corner is the z-location within the volume. (A) Membranes of the degradation compartment segmented (red) and content (blue). Scale bar: 2 µm. (B) Density (ratio of the content volume within the degradative compartment) as a function of the volume of the degradative compartment showing the presence of four unrelated populations (quadrant count ratio = 0.037). (C-E) Enlarged panels from the coloured boxes in (A), highlighting the membrane contact sites between the degradation compartment (DC) or empty vesicles and other organelles including mitochondria (m) and chloroplast (C) or an absence of contact with, for example, the Golgi apparatus (GA). The limits of these contacts are indicated by a black arrow. Scale bars: 500 nm (C), 250 nm (D-E). (F-G) Enlarged panels of the eyespot and 3D segmentation, proximal to the reservoir (R). Compartments in proximity (V: vesicle, C: chloroplast). Scale bar: 250 nm.

### Classifying the neuronal network of an adult mouse brain

Finally, one of the major challenges in volume cryo-SEM imaging is the imaging of tissues, among which brain, rich in myelin, remain a challenge. Indeed, serial SEM analysis of brain is often limited to young individuals where myelinisation is not fully established^19,37,40,58^. Moreover, the ability to study the thickness of the myelin in the case of normal or pathological development has been limited by artefacts due to the fixation, dehydration and staining required for room temperature analysis^59^ or require specialized sample preparation^60^. However, by modifying the scanning pattern, we have observed the myelinated structure in an adult brain cortex in a near-native state (Extended Data Figure 10). We observed axons with thick or thin myelination (Ax or ax respectively) in all areas within the field of view, including areas where the axons were bundled. The reduction of charging artefacts also allowed the observation of structure within the axon (inner tongue of the oligodendrocyte, mitochondria, vesicle) or in their vicinity (mitochondria, vesicles, ER).

The acquisition of a volumetric dataset using interleaved scanning and a short dwell time allowed us to understand the myelination organisation (Supplementary movie 2). In the central nervous system (CNS), myelin is formed by oligodendrocyte cells wrapping around the axon. It has been shown that the wrapping of the myelin sheath is a combination of two coordinated events: the wrapping of the leading edge of the inner tongue (the part of the oligodendrocyte closest to the axon) and the lateral extension of myelin membrane layers towards the nodal region (the part of the oligodendrocyte the furthest from the axon)^40^. This growth is supported by a system of cytoplasmic channels providing shortcuts for the transport of membranes. Previous work established this process using the optic nerve of young mice (P10-60), which was dissected, HPF and then freeze substituted and stained. We were able to reproduce and extend this work in non-denaturing conditions in an adult mouse cortex by distinguishing thin and thick myelinated axons. We defined and tracked thin and thick myelinated axons in 3 dimensions (Figure 5A). Thin myelinated axons have myelin sheaths of 19 ± 4 nm, while thick myelinated axon present a sheath of 40 ± 11 nm (Figure 5 and Extended Data Figure 10). The thin myelinated axon population is the only type for which we were to observe some axons along their entire length within our volume (1134 µm^3^) (Figure 5 yellow box and in A, B and D). For this population we also observed the network of cytoplasmic channels across a 1.6 ± 1.3 µm^3^ volume inside the myelin sheaths (Figure 5 A and D).

**Figure 5:**
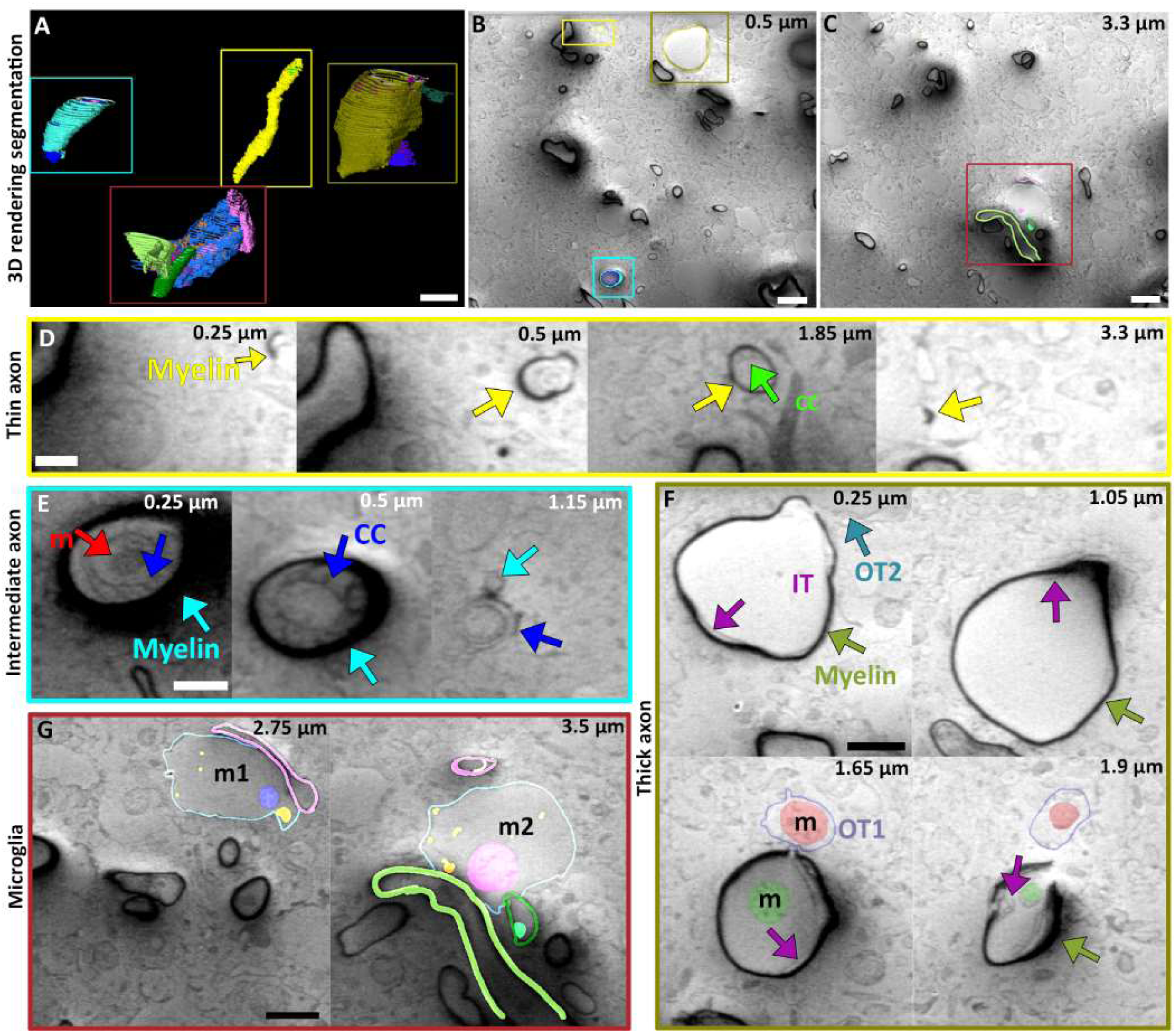
SEM volume imaging of mouse brain. A 118-day old mouse brain imaged at 90° with respect to the FIB milled sample plane using 100 ns dwell time x100 repetitions. A 1334 µm^3^ volume in focus was aligned and manually segmented for the region of interest. (A) segmented volumes. (B and C) slices from the volume with overlayed segmentation. Scale bar 2 µm. Boxes in A are superimposed on B and C and enlarged in boxes (D-G) with the respective coloured box containing representative slices in the volume. Numbers in one of the corners are the z-location within the volume. (D) thin myelin axon with internal cytoplasmic channels (CC) in green. Scale bar: 250 nm. (E) intermediate axon with extended CC, mitochondria (m). Scale bar: 250 nm. (F) thick myelin and large axon and the contact with 2 oligodendrocytes showing the inner tongues (IT) and the outer tongs (OT). Scale bar: 500 nm. Mitochondria are also visible in the axon and one of the outer tongues. (G) microglia cells and contacts with the different axons showing extended contact between the different cells, as well as internal organelles including mitochondria (m), vesicles (v) and inner tongues (IT). Scale bar 500 nm.

Within the thick myelinated axon population, axons with diameters under or over 1 µm (Figure 5 and Extended Data Figure 10) were observed, which we as denote intermediate and large axons respectively. The intermediate axon population shows an extended network of cytoplasmic channels (6.6 ± 2.1 µm^3^) to support the extended growth of the axon (Figure 5A, B and E). On these axons, we also observed the progression from thin to thick myelin as the myelin compacts (example Figure 5E).

In some cases, it was possible to identify unique features in the data where the neurons changed morphology along their length. For example, we observed the start of a large axon (∼1.5 µm in diameter) (Figure 5A, C and F) where, in this instance, we could not observe the cytoplasmic channels, as had been observed in other examples. Instead, breaks were visible within the myelin sheath associated with membranes from other cells, most probably oligodendrocytes, which are associated with additional membranes under the existing myelin. This suggests that large axons with thick myelin support myelination by the constant addition of oligodendrocytes tongues, in contrast to the mechanism of myelination in thin or intermediate axon populations.

The reduction of charging artefacts also allowed the visualisation of the immediate axon environment, identifying the extensive contacts that microglia have with the myelinated axon. Microglia cells play a crucial role in the maintenance of the myelin^61^. We observed that a single microglia cell has 3 extended contacts with 3 different axons (Figure 5A, C and G). At this contact the plasma membrane of the microglia is sometimes impossible to distinguish from the myelin sheath, in agreement with their role in immunological survey of the CNS^62^.

Using our modified scanning pattern, we observed and characterised different axons based on their diameter and myelin thickness which in turn supports a different strategy in the formation and maintenance of the myelin sheath. The ability to observe different axonal phenotypes could be critical when mapping the different functions of axons within the brain, where some axons carry messages within the CNS while others to the periphery and some ensure anterograde while other retrograde signalling. We also observed the interaction of microglia cells with different axons, demonstrating that absence of charging artefacts allows for more features to be identified inside and outside the axons.

## Conclusions

Volume imaging using FIB/SEM is a well-established technique used in both materials and life sciences^3,63–65^. However, most work reported in the literature describes results from samples at room temperature, which in the life sciences requires sample preparation that may introduce artefacts ^66,67^. At cryogenic temperatures, the data can also be corrupted with curtaining or charging artefacts^19,26,27,58^. Here we have demonstrated that modifying the scanning pattern reduces the charging artefacts as illustrated for different exemplars of vitrified biological samples. This paper focuses solely on an interleaved 3×3 scan in a frame size of 2048×2048 pixels. However, the relation between the interleaved steps and the image resolution requires further examination with regards to the probe size and spatial separation between scan positions. In principle, any number of pixels in the 2D array can be skipped while scanning in the x and y directions and alternative irregular scan geometries are also possible^31^. However, as the scan step increases, the response of the scan deflectors may introduce errors in the probe positioning, leading to image distortions^68^.

In this work, we have demonstrated that skipping a small number of pixels allows charge dissipation while maintaining low image distortions. However, we also observed that there is further scope to optimise the interleaved parameters as on some occasions charging artefacts, appearing as halo surrounding a charging centre, which is particularly noticeable around LDs (Figure 3F or I) or thick myelinated axons (Figure 5 F). Since the charge and discharge rate depends on the electrical resistivity and surface area of the sample ^12^, the optimal number of pixels to skip is to a large extend dictated by the sample itself.

To demonstrate the impact of interleaved scanning on the reduction of charging artefacts, the number of parameters (accelerating voltage, imaging angle, beam current, pixel size, total fluence, working distance) have been deliberately restricted in this work. However, this may lead to non-optimal image acquisition. For example, when imaging normal to the milled sample plane the energy crossover will change^69^, and if not compensated this will introduce additional charging artefacts (Extended Data Figures 6-7). Similarly, HPF samples have reduced charge dissipation capacity as they form of a solid layer of non-conductive material and often require a decreased accelerating voltage for imaging^19^.

In our experimental set up, we sometimes observed ‘burn spots’ (approximately 100 nm size) that are not related to a specific biological structure but arise from beam damage. These are present when using the external scan engine and are more obvious during interleaved scanning (Figure 3C, I and Extended Data Figure 10E, F) due to the scan being paused and the beam left unblanked between acquisitions. This pause occurs during the transfer of the analog signal to the host PC in the hardware used and during the data type conversion, both of which occur in millisecond time scales.

As volumetric SEM imaging is used to track structures of interest, segmentation is frequently applied to the acquired images. To measure the impact of the reduction of charging artefacts on image interpretation, we applied the Segment Anything Model to our dataset. The presence of charging artefacts led to the incorrect identification of non-existent or distorted structures ( Extended Data Figure 8). Conversely, the systematic absence of artefacts allowed more features to be selected with increased complexity. This improvement in segmentation due to the reduction of charging artefact minimises the human intervention requiremed, making on-the-fly segmentation and feature recognition for automated SEM targeted milling feasible in the future. To demonstrate the utility of our modified scanning approach we have imaged several different relevant biological structures. LD, DC and axons were chosen as their compositions (mostly lipids and/or metabolites) and their insulating nature make them extremely sensitive to sample preparation artefacts and these structures are a known source of charging artefacts. Imaging these structures at high resolution is therefore at the limit of state-of-the-art SEM. Here we observed different type of organelles’ MCS as a results of being able to image subtle differences allowing us to decipher membrane tethered or hemi fused in LD/ER MCS in mammalian cells. In *E.gracilis*, we were able to segment the network of DC, identify their different sub-populations and hypothesise that MCS between DC and other organelles, such as mitochondria and chloroplasts, leads to the emptying of the DC content. Better identification of features, resulting from modification of the scanning pattern could inform additional correlative studies ^19,27,28,70,71^. Finally, we analysed two axons with different myelin thicknesses in adult mouse cortex. We imaged perpendicular to the milled plane of the sample to increase contrast^19^ and due to the absence of artefacts were able to map specific axon phenotypes. These different phenotypes could not be identified at room temperature due to sample preparation and/or imaging artefacts.

In summary, we have demonstrated that by modifying the scan pattern we can reduce charging artefacts, removings constraints on sample preparation for imaging specimen of varying conductivity. This effectively inverts the paradigm that samples should adapt to the constraints of the microscope and it is now possible for the imaging system to adapt to the sample specificity. Combined with automated segmentation of features, volume SEM imaging with reduced artefacts provides a tool for automated recognition in FIB lamella preparation for *in-situ* structural biology, creating a workflow with the potential to impact life sciences and medical diagnostics.

## Materials and Methods

### Sample preparation

RPE-1 (CRL-4000) cells were obtained from the ATTC and cultured as per the supplier recommendation. For plunge freezing the cells were grown on EM grids (gold/200mesh/R2.2 either UltrAufoil or Quantifoil) before vitrification using a Vitrobot (ThermoFisher Scientific) as previously reported^19^.

A 118-days-old (C57BL-6) mouse was euthanized following a schedule 1 procedure via intraperitoneal injection of sodium pentobarbital followed by decapitation following licensed procedures approved by the Mary Lyon Centre and the Home Office UK. The cortex was prepared as previously reported^19^.

*Euglena gracilis* Klebs CCAP 1224/5Z was routinely cultured in *E. gracilis* medium and Jaworski’s medium mixed in 1:1 proportions^72^. Cultures were maintained at 15°C under a 12:12h light:dark regime. Illumination was provided by cool white LED lamps with a photon flux density of 50 µmol.m^-2^.s^-1^ at the surface of the culture vessel. For plunge freezing the cells were pipetted (4 µl) on EM grids (gold/200mesh/R2.2 either UltrAufoil or Quantifoil) in a humidified chamber (98% RH), single side back blotted (20 s) before vitrification by plunging into liquid ethane using an EM GP Plunge Freezer (Leica microsystems, Austria).

### SEM imaging

An external scan engine (Quantum Detectors) was interfaced to a Helios Hydra G5 (ThermoFisher Scientific) using the available lithography connection. The cryo-stage and the anti-contamination strategy is adapted from the Aquilos II system (ThermoFisher Scientific). The scan inputs to control the SEM deflectors were controlled by the Quantum Detector scan engine and the signal from the scintillator/photomultiplier secondary electron detector was connected to one of its analog inputs. The scan engine software is based on a Python module with the data saved as 2D NumPy arrays, as int16 data type by default. The pattern for the interleaved scan was generated using an in-house Python script (see data and code availability) and skips two pixels in the x scan direction and two lines in the y scan direction, as illustrated in Figure 1. This script provides a text file of x and y coordinates, which is read by the external scan generator to control the position of the electron probe for imaging. The control is only used for the SEM probe deflectors. AutoScript (ThermoFisher Python-based scripting software) was used to integrate SEM imaging using the programmable scan engine during FIB/SEM volumetric imaging. To limit the parameter space explored, we defined the base parameters to match our previous work^19^. In summary, the parameters used for SEM imaging were; accelerating voltage 1.2 kV, beam current 6.3 pA, probe size approximately 2 nm at FWHM, dwell time 100 - 1000 ns, 10 – 100 repetitions, immersion mode (suction tube 70 eV), detector: in-lens for secondary electrons, pixel size 6.34 nm. The detector voltage offset and gain were maintained per dataset but not between samples of the same sample. We did not apply flyback delay during acquisitions using frame integration for both raster and interleaved patterns, to ensure that all images were acquired with the same electron fluence. This flyback delay is commonly applied at the start of each raster scan line to wait for stabilization of the probe after it is moved from the end of one line to the next one^68,73^. However, this flyback delay was applied to raster line integration acquisitions as these were acquired using the inbuilt functionality of the microscope (accessed through the microscope’s user interface) which does not allow deactivation of the flyback delay of (approximately 70 µs for the conditions used in our experiments). Beam damage or charging effects were not observed on the left hand-side of line integration acquisitions since the probe is only stationary for a very short time. The acquisitions were cropped to perform the analysis on images without distortions, as described below. For consistency in comparisons, the raster line integration images were also cropped.

### Automation with SerialFIB

As for previous implementations of customized SEM imaging procedures^19^, the scanning strategies reported here were integrated to the software package SerialFIB^74^. The volume imaging script developed for different milling and image acquisition positions that enables perpendicular SEM imaging was extended using the scripts above to control the scanning patterns. In order to compensate an issue when switching to the SEM scanning after FIB milling, a preview image at a dwell time of 25ns was acquired prior to the interleaved scanning exposure. The data acquisition was implemented using the customized scripts described above, writing out single frames per interleaved scan after acquisition.

### Pre-processing

#### Estimation of the spatial extent of flyback distortions

The absence of a flyback delay leads to distortions and poor image quality on the left side of the acquisitions. For this reason, to remove distortions, raster and interleaved acquisitions were cropped. This number of pixels was estimated from the interleaved images acquired using the shortest dwell time since these settings will result in maximum distortions. This factor is considered as a figure of merit for all the acquisitions, both raster and interleaved.

We implemented an automated approach (data and code availability section), to statistically estimate this factor by selecting a small region in each frame of a sequential acquisition that shows obvious flyback distortions. The same region is selected for all frames in the experiment, as shown in Supplementary Figure 1 A Gaussian filter, σ = 7 pixels, was applied to increase the SNR and subsequently a Sobel filter (3×3 kernel), for gradient approximation and a threshold for edge detection and to convert the grey-scale images into binary images. The result is a vertical band whose right edge (rightmost pixel with intensity value one) represents the spatial extent of the flyback distortions. This approach was applied to the RPE-1 dataset acquired using 100 ns dwell time, 100 repetitions. A histogram was calculated from the measurements of all the acquisitions, and the mean value of the normal distribution taken as the figure of merit. The histogram indicates 64 pixels as the extent of the distortions. We considered a valid approximation since the Gaussian filter smooths fine details, inherently extending the edge of the apparent flyback distortions. The outliers on the histogram are the result of the edge detection process failing to find clear edges in a noisy background.

#### Subframe preparation

Since a relatively large amount of data was collected, particularly during volumetric imaging, the 2D Numpy arrays were converted to a uint8 data type during acquisition and stored as raw data; given that grayscale levels of 8-bit images are not visually discernible and are adequate for the image processing performed in this work^75^, as demonstrated during nuclei instance segmentation^76^, for instance. The conversion was done by shifting the histogram of the acquisitions to a positive range and subsequently re-scaling the pixel intensities. Immediately after acquisition, the 2D Numpy arrays were then grouped and stacked to create a 3D array and saved in a tiff format (see data and code availability section). Cropping was performed post-acquisition to remove flyback artefacts.

#### Subframe to frame (image)

For frame integration, the frames were aligned using MotionCor2^25^, with 5×5 patch and 20% overlap.

#### Frames to stack (movie)

For volumetric data, the frames (or images) were aligned using a SIFT^77^ plugin in Fiji^78^ using the default parameters for the affine option without interpolation^77^.

### Histogram analysis

The mean value of the histograms of aligned subframes were calculated using the Numpy library in Python. Mean values were calculated from the cropped data (1920×1920 pixels) using a script for batch processing in Fiji.

### Myelin sheath thickness measurement

Using the volume of the brain acquired, lines were manually placed on the myelin sheath where intense dark pixels were present. We used 10 measurements over 3 independent axons and measured their length using the Fiji build-in measurement tool. T-tests were performed over the two populations that show that they are independent with a confidence of 1%.

### Segmentation and subsequent data analysis

Aligned and cropped subframes, forming a frame or image, were filtered using a Fiji mean filter (using a 3×3 filter kernel).

#### Unsupervised SAM segmentation

Images were loaded using a Python script and processed by the SamAutomaticMaskGenerator from Segment Anything^51^. The Segment Anything model used was ViT-H with 32 points_per_side, a value of 0.92 for pred_iou_thresh and 0.9 for stability_score_thresh with other parameters set to default values. The perimeter and area of each segmented component was calculated with skimage regionprops and a ‘complexity’ score was calculated as the squared perimeter of the component divided by the area. These scores were summed for all components in the segmentation to produce a final image complexity score.

#### Manual segmentation

For *E.gracilis,* in addition to the mean filter, the ‘remove background’ option ^79^ in Fiji was used with a using a rolling ball of 50 pixels and the light background option.

Amira (ThermoFisher Scientific) was used to manually segment volumes which were then exported as a tiff stacks for further analysis using Fiji. BioVoxxel 3D and Image 3D suite and their dependencies were used for volume calculations of the different segmented objects^80–82^.

## Supporting information

Supplementary figure 1, supplementary table 1 and 2 and legend for the supplementary movies 1 and 2

Extended figure 1 to 10 with their respective legend

## Acknowledgments

The authors would like to acknowledge the help of Matthew Case for analysis of the brain samples. Kamlesh Patel from Thermosfisher Scientific, Leo Jay-Black and Ben Bradnick from Quantum Detectors for the integration of the scan engine/microscope and for the Python module help, respectively. Elaine Ho for her help with Motioncor2. This work was supported by the Wellcome Trust through the Electrifying Life Science grant (220526/Z/20/Z to JHN) and a Wellcome Trust Development Award (225902/Z/22/Z to MG). The Rosalind Franklin Institute, funded by the UK Research and Innovation, Engineering and Physical Sciences Research Council.

## Author contributions

AV, conceptualized and developed the acquisition protocol, acquired. TG integrated the scan engine, acquired data. SK automatized the scan engine protocol. AP performed the automated segmentation. JZ performed manual segmentation. JS, CG and MK prepared the sample, WB performed preliminary experiment MK prepared samples, RAF, JHN, JSK, MCD, MG, AIK supervised the work. MD also performed manual segmentation and interpreted the data. All authors commented on and edited the manuscript.

## Data and code availability

Scripts for data acquisition and pre-processing are available on the Rosalind Franklin Github (https://github.com/rosalindfranklininstitute/fib-sem-charge-mitigation). For SerialFIB the software and specific scripts for the scan engine are available on GitHub (https://github.com/sklumpe/SerialFIB). The volume acquired for *E.gracilis* and adult mouse cortex, as well as the segmented volumes, are available on EMPIAR (47486076 and 47486077).

