## Supplementary figure 1, supplementary table 1 and 2 and legend for the supplementary movies 1 and 2 for "Reduction of SEM charging artefacts in native cryogenic biological samples": Velazco et al supp info.pdf

### Supplementary information:

| | | Raster-line integration | | | Raster-frame integration | | | Interleaved-frame integration | | |
|---|---|---|---|---|---|---|---|---|---|---|
| | | 100x100 | 500x20 | 10x1000 | 100x100 | 500x20 | 10x1000 | 100x100 | 500x20 | 10x1000 |
| Raster-line integration | 100x100 | | 61.7% | 15.9% | 90.8% | 18.1% | 10.7% | 2.4% | 90.2% | 89.6% |
| | 500x20 | | | 22.9% | 54.7% | 10.3% | 6.7% | 9.5% | 52.7% | 58.6% |
| | 10x1000 | | | | 13.7% | 34.7% | 1.8% | 57.1% | 12.7% | 17.4% |
| Raster-frame integration | 100x100 | | | | | 21.5% | 13.1% | 1.9% | 99.9% | 97.0% |
| | 500x20 | | | | | | 78.0% | 0.1% | 18.2% | 34.7% |
| | 10x1000 | | | | | | | 0.1% | 10.1% | 25.4% |
| Interleaved-frame integration | 100x100 | | | | | | | | 1.4% | 4.1% |
| | 500x20 | | | | | | | | | 96.9% |
| | 10x1000 | | | | | | | | | |

**Supplementary table 1: T test analysis of different samples imaged using different scanning strategies and electron fluence**. Green values represent statistical differences between the populations (2 tails, difference variance).

| | | Raster-line integration | | | Raster-frame integration | | | Interleaved-frame integration | | |
|---|---|---|---|---|---|---|---|---|---|---|
| | | 100x100 | 500x20 | 10x1000 | 100x100 | 500x20 | 10x1000 | 100x100 | 500x20 | 10x1000 |
| Raster-line integration | 100x100 | | 51.3% | 44.8% | 6.2% | 31.8% | 29.6% | 2.6% | 22.8% | 38.8% |
| | 500x20 | | | 89.8% | 85.0% | 85.1% | 80.5% | 36.9% | 95.0% | 93.3% |
| | 10x1000 | | | | 98.9% | 96.6% | 92.0% | 47.6% | 92.4% | 81.9% |
| Raster-frame integration | 100x100 | | | | | 96.4% | 89.6% | 20.2% | 82.8% | 66.6% |
| | 500x20 | | | | | | 94.5% | 41.6% | 86.1% | 74.0% |
| | 10x1000 | | | | | | | 46.9% | 80.1% | 68.9% |
| Interleaved-frame integration | 100x100 | | | | | | | | 21.2% | 18.3% |
| | 500x20 | | | | | | | | | 83.3% |
| | 10x1000 | | | | | | | | | |

**Supplementary table 2: T test of different samples imaged using varied scanning strategy and fluence**. Green values represent statistically significant differences between the population (2 tails, difference variance).

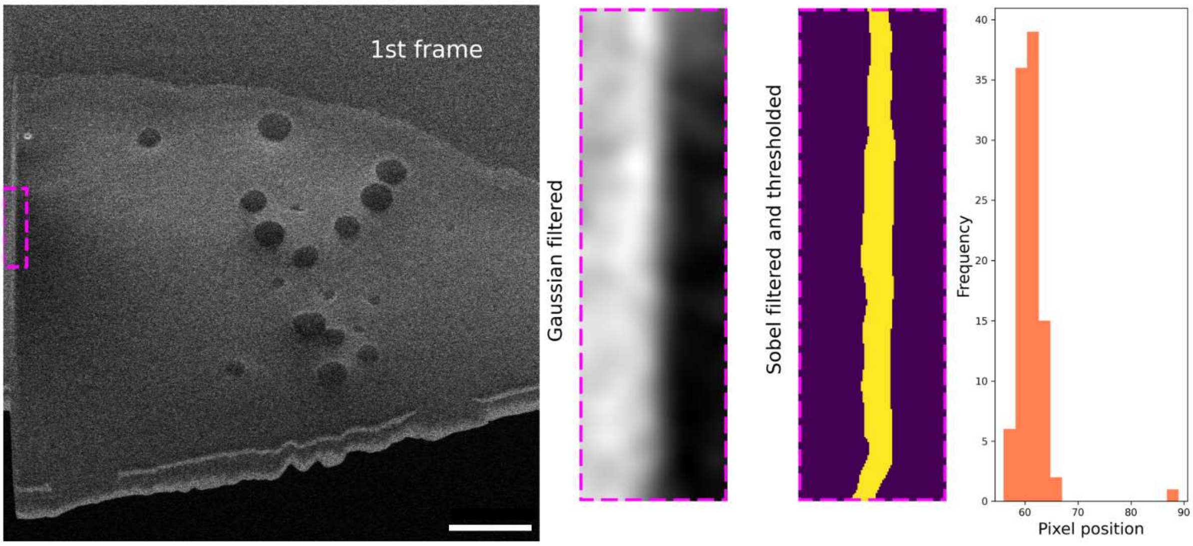


**Supplementary figure 1: Estimating the spatial extent of flyback distortions.**
Processing of images from the vitrified RPE-1 sample, acquired using an interleaved scan, 100 ns dwell time x100 repetitions. Area showing distortions is indicated by a colored box in the image on the left. The same area was processed for all the frames as described in the text. Scale bar: 2 µm.

**Supplementary movie 1: Volume acquisition of vitrified *E.gracilis*.** Vitrified *E.gracilis* imaged at 52° with respect to the FIB milled sample plane using 100 ns dwell time x100 repetitions. A 1600 µm³ volume in focus was aligned and manually segmented for region of interest.
**Supplementary movie 2: Volume acquisiton of vitrified mouse brain cortex**. A 118-day old mice corte imaged at 90° with respect to the FIB milled sample plane using 100 ns dwell time x100 repetitions. A 1334 µm³ volume in focus was aligned and manually segmented for region of interest.

.
