## Extended figure 1 to 10 with their respective legend for "Reduction of SEM charging artefacts in native cryogenic biological samples": Velazco et al extended data.pdf

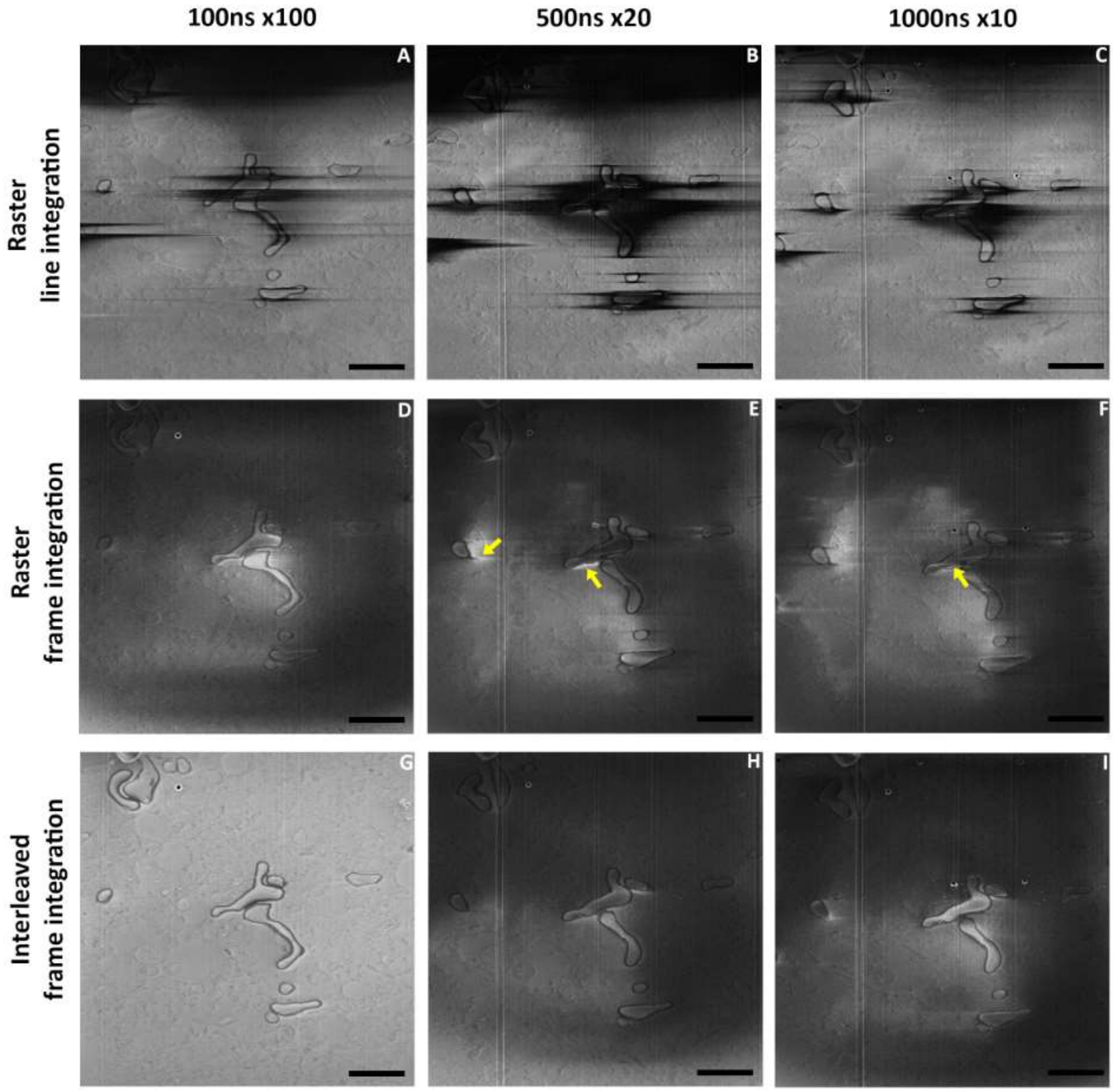


**Extended Data Figure 1: Effect of electron fluence distribution on charging artefacts (SEM imaging at 52º angle to the FIB milled sample plane).**
(A-I) Representative images of brain tissue for different pixel fluences and raster or interleaved scan pattern with (G) interleaved frame integration at 100 ns x 100 showing the greatest improvement in charging artifacts. Arrows: bright charging artefact. Dashed rectangle: improved region with higher fluence. Scale bar: 2 µm.

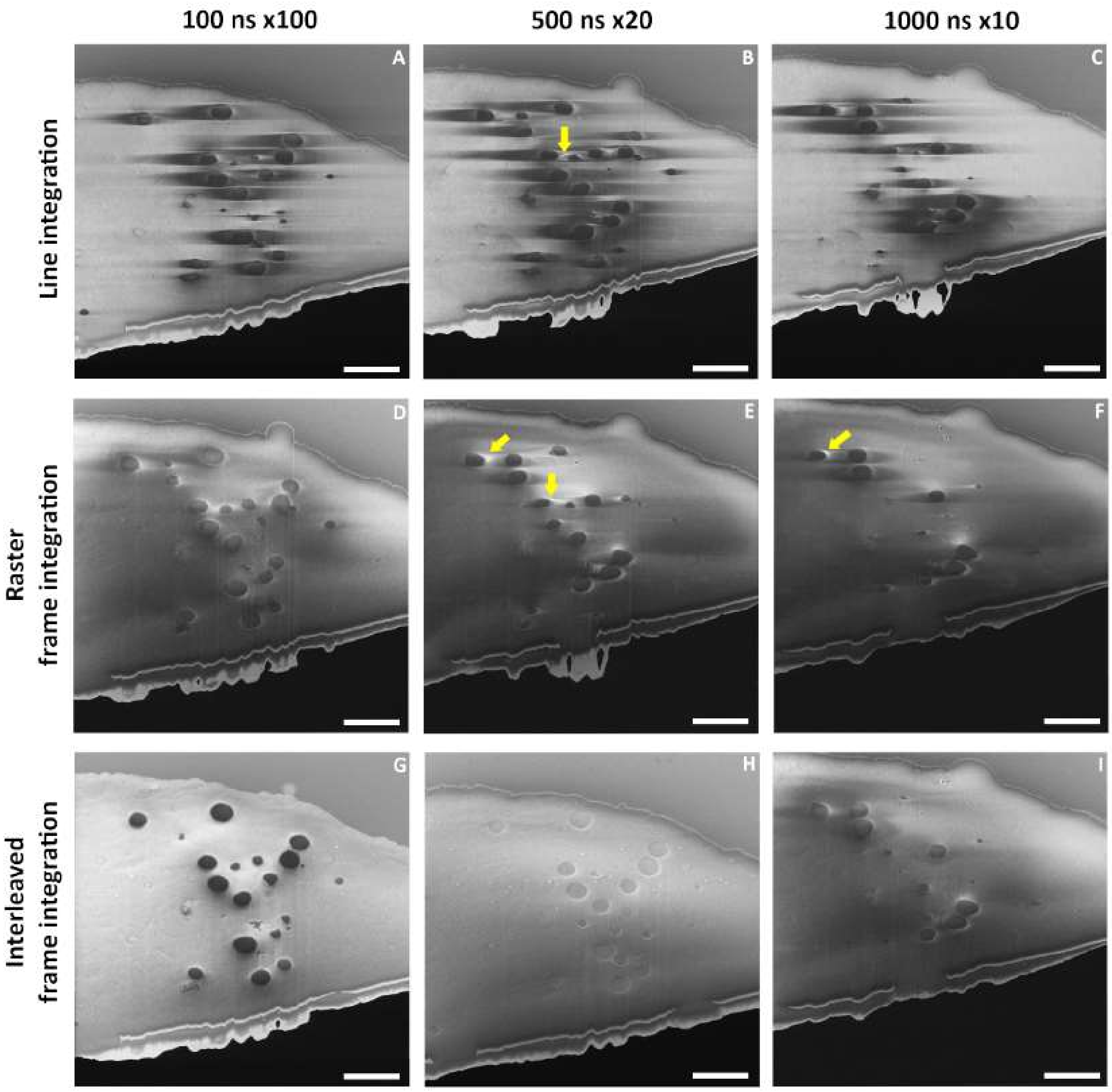


**Extended Data Figure 2: Effect of electron fluence distribution on charging artefacts.**
Vitrified RPE-1 cells imaged at 52° with respect to the FIB milled sample plane using the same parameters for dwell time and number of repetitions (for frame or line integration) but different scanning strategies. To avoid accumulation of potential beam damage, the area was milled between image acquisitions (50 nm steps). Arrows indicate the presence of bright charging artefacts. Scale bar: 2 μm.

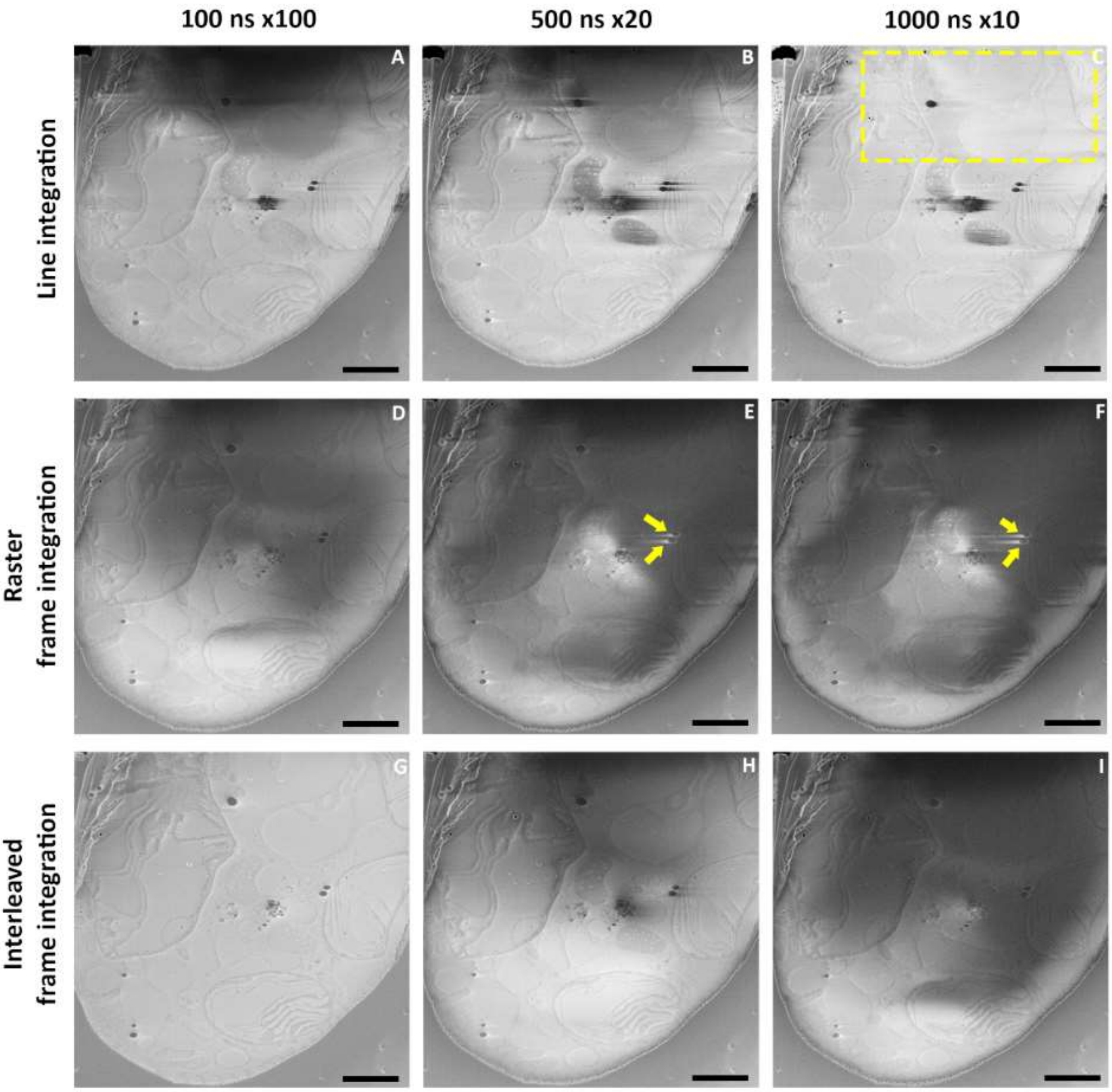


**Extended Data Figure 3: Effect of electron fluence distribution on charging artefacts.**
Vitrified *E. gracilis* imaged at 52° with respect to the FIB milled sample plane using the same parameters for dwell time and number of repetitions (for frame or line integration) but different scanning strategies. To avoid accumulation of potential beam damage, the area was milled between image acquisitions (50 nm steps). Arrows indicate the presence of bright charging. Dotted rectangle: improved area with higher fluence strategy. Scale bar: 2 µm.

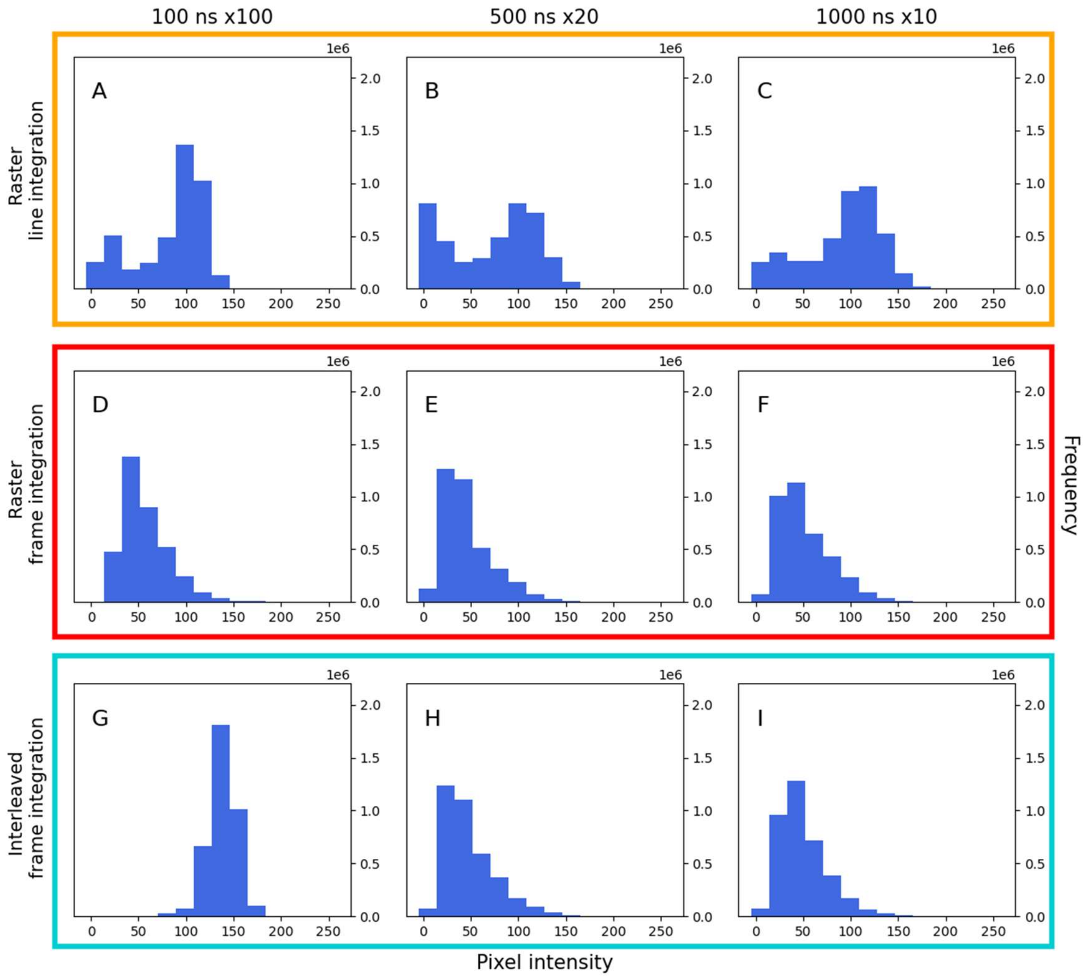


**Extended Data Figure 4: Histograms from images of brain tissue acquired at different electron fluence.**
(A-I) Histograms of representative images of brain tissue for different pixel fluence strategies and raster or interleaved scan patterns. To avoid accumulation of surface beam damage, the area is FIB milled between image acquisitions by 50 nm.

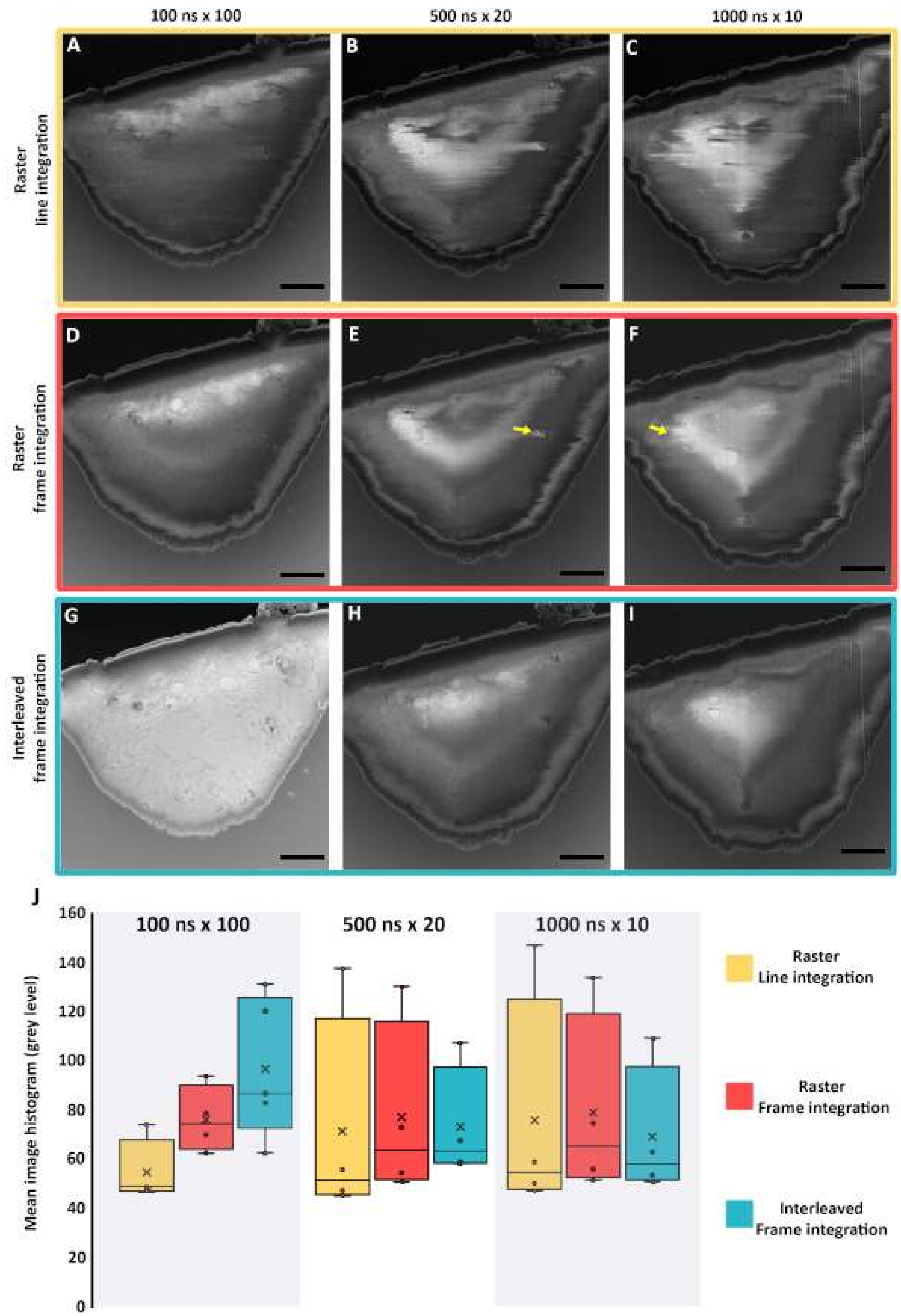
100 ns x 100
500 ns x 20
1000 ns x 10
Raster
line integration
Raster
frame integration
Interleaved
frame integration
A
B
C
D
E
F
G
H
I
J
Mean image histogram (grey level)
0
20
40
60
80
100
120
140
160
Raster
Line integration
Raster
Frame integration
Interleaved
Frame integration

**Extended Data Figure 5: Effect of electron fluence distribution on charging artefacts (SEM imaging perpendicular to the FIB milled sample plane).**

(A-I) Images of RPE-1 cell for different pixel fluence strategies and raster or interleaved scan patterns with (G) interleaved frame integration at short 100 ns x 100 showing the greatest improvement in charging artifacts. Arrows: bright charging artefact. Dashed rectangle: improved region with higher fluence strategy. Scale bar: 2 µm. (J): Mean of image histograms for different populations (n>4) from vitrified RPE-1, *E.gracilis* and mouse brain represented as a circle. Population median (middle line), population mean (cross), median of the 1$^{st}$ quartile of the population (bottom line of the box), median of the 3$^{rd}$ quartile of the population (top line of the box). Vertical lines extend to minimum and maximum values. Statistical analysis is given in Supplementary Table 2. Images acquired using interleaved scanning with frame integration associated with short dwell time and high integration (100 ns x100) form a dataset for which the mean of the histogram is at 97, closest to 127 which is the mid-range histogram intensity value of 8-bit images (0-255 range).

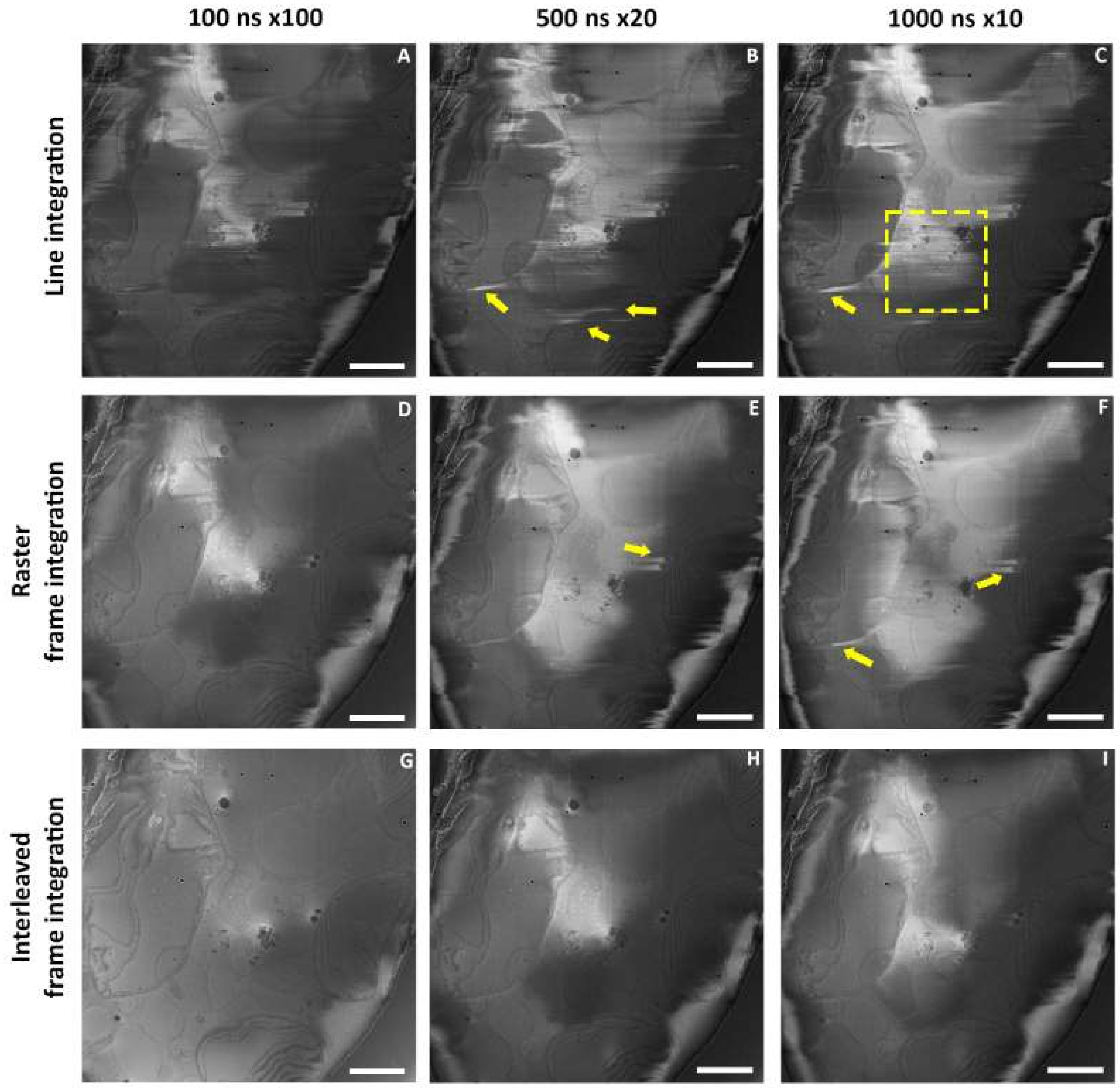


**Extended Data Figure 6: Effect of electron fluence distribution on charging artefacts.**
Vitrified *E. gracilis* imaged at 90° with respect to the FIB milled sample plane using the same parameters of dwell time and number of repetitions (for frame or line integration) but different scanning strategies. To avoid accumulation of potential beam damage, the area is milled between image acquisitions (50 nm steps). Dotted rectangle: improved region with higher fluence. Arrows: bright charging artefacts. Scale bar: 2 µm.

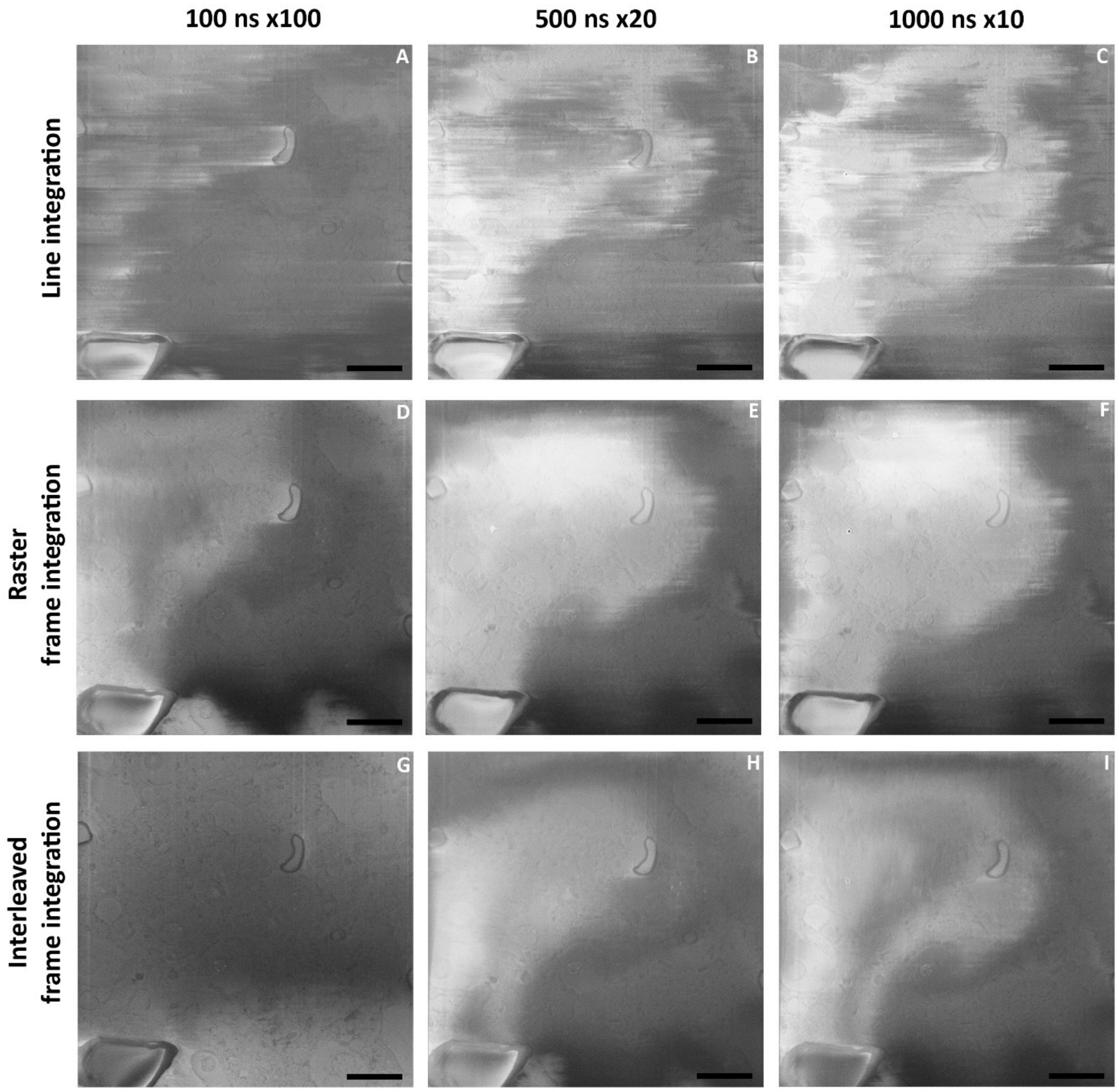


**Extended Data Figure 7: Effect of electron fluence distribution on charging artefacts.**
Vitrified 118-days old mouse brain slice imaged at 90° with respect to the FIB milled sample plane using the same parameters of dwell time and number of repetitions (for frame or line integration) but different scanning strategies. To avoid accumulation of potential beam damage, the area was milled between image acquisitions (50 nm steps). Scale bar: 2 µm.

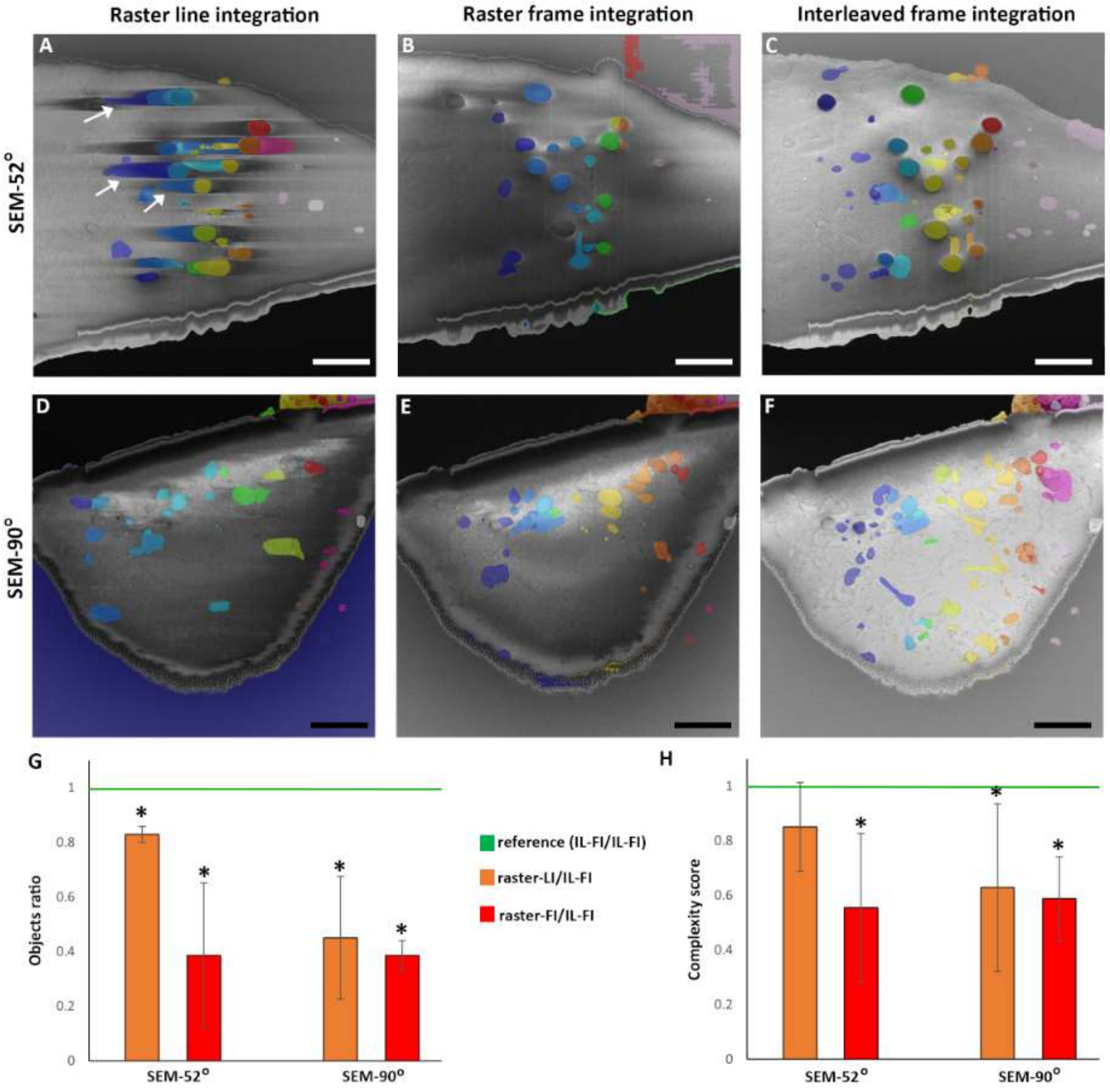


**Extended Data Figure 8: Effect of scanning strategy on segmentation output.**

Vitrified RPE-1 cells imaged at 52° (A to C) and 90° (D to F) with respect to the FIB milled sample plane using 100 ns dwell time x100 repetitions, raster line integration (A, D), raster frame integration (B, E) and interleaved frame integration (C, F). (A-F) SEM images (grey) overlaid with segmented areas (rainbow). White arrows indicate segmentation outputs at locations corresponding to charging artefacts. Scale bar: 2 µm. (G and H) analysis of vitrified mouse brain, RPE-1 and *E.gracilis* samples, * indicate statistical difference ($p<5\%$). Ratio of number (G) or complexity (squared perimeter divided by the area) (H) of objects detected using Segment Anything Model. Parameters were normalized for all datasets. The number of objects picked were not manually corrected to objects which were not biological features. For example, where a picked object is entirely composed of charging artefacts it was not curated. Values obtained from raster line integration (LI) or raster frame integration (FI) acquisitions were compared to interleaved frame integration (IL-FI) acquisitions.

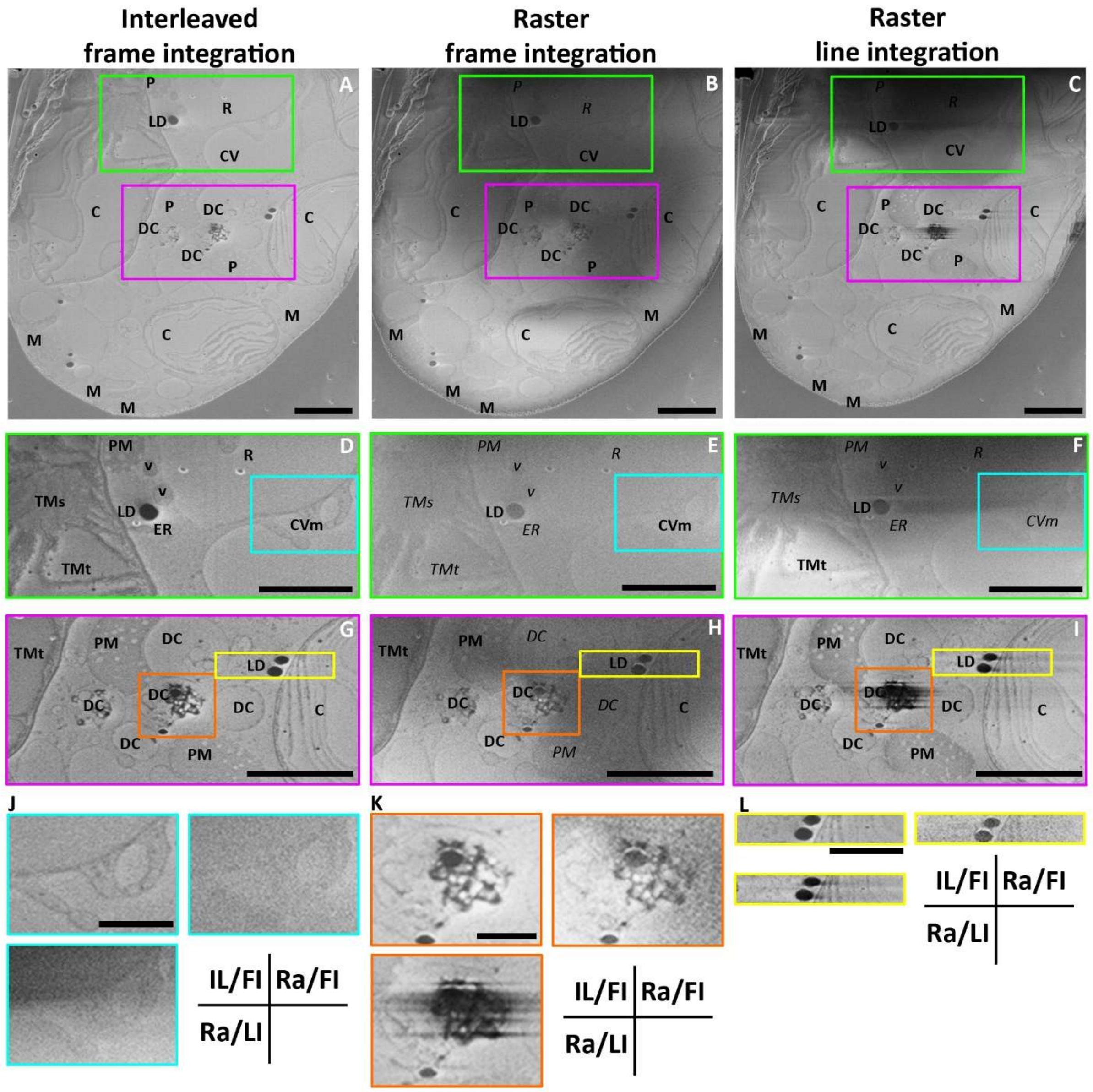


**Extended Data Figure 9: Charging artefact mitigation in vitrified *E.gracilis* allows the observation of different vacuoles and degradation compartments.**

Images of *E.gracilis* cells at 52° with respect to the FIB milled sample plane, 100 ns dwell time x100 repetitions using (A) raster line integration (Ra/LI) (B) raster frame integration (Ra/FI) (C) interleaved frame integration (IL/FI). P: paramylon, R: reservoir, CV(m): contractile vacuole (membrane), LD: lipid droplet, C: chloroplast, DC: degradative compartment, M: mitochondria, v: vesicles, ER: endoplasmic reticulum, TMs/t: thylakoid membrane sheet/tubes. Labels in italic indicate structures that could not be observed. (D to L) color-coded highlighted areas. Interleaved frame integration allows identification of compartments otherwise hidden by charging artefacts. (D-L) Brightness and contrast were optimized to improve visualisation. (D to F), enlarged areas showing the location of the contractile vacuole and the reservoir, panel in J shows increased contrast for IL/FI. (G to I and K) highlights of the modified scanning strategy enables observation of the content and the contact points of the degradation vesicle which are otherwise partially/totally obscured. (L) highlights that the presence of putative lipid droplets is not associated with the presence of charging artefacts allowing the observation of these compartments and their immediate surroundings. A-I scale bars: 2 μm, J-K scale bars: 1 μm.

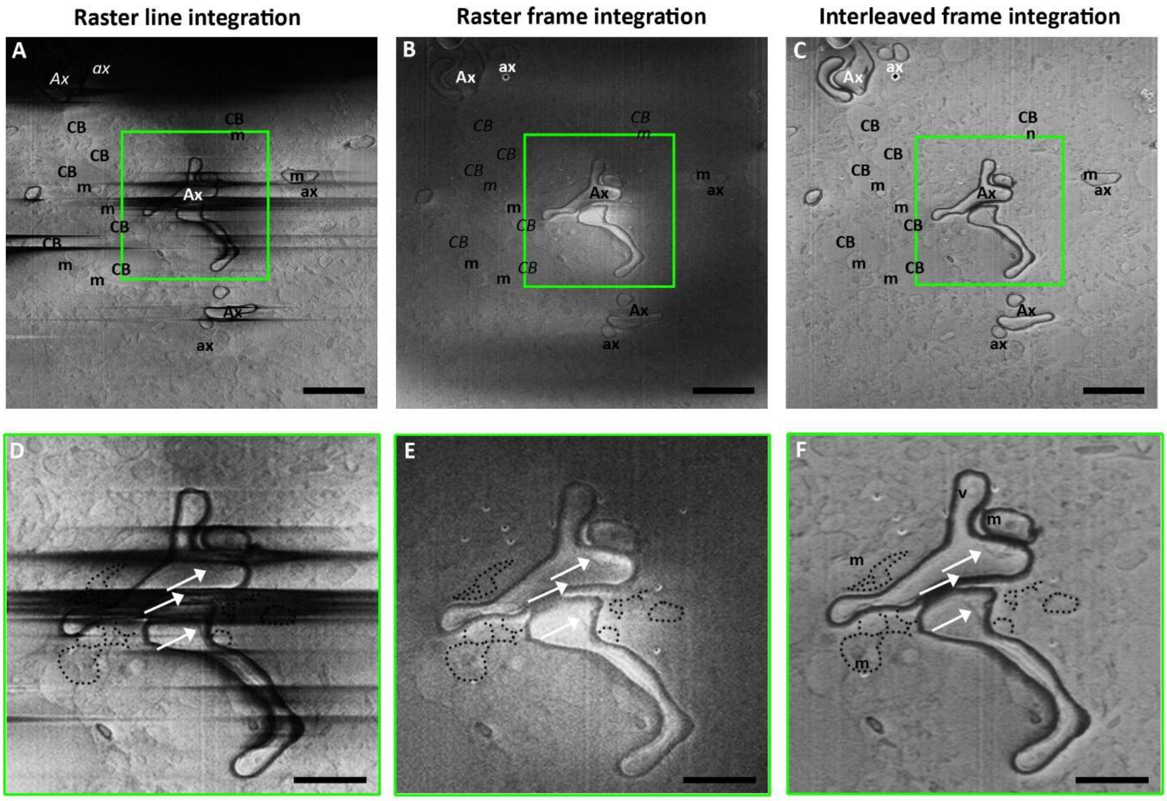


**Extended Data Figure 10: Charge mitigation in SEM imaging of mouse brain tissue allows imaging of different myelinization thickness and axon content.**

A 118-day old mice brain imaged at 52° with respect to the FIB milled sample plane using 100 ns dwell time x100 repetitions. Brightness and contrast were optimized for visualisation. (A-C) CB: cellular body, Ax: thick myelin, ax: thin myelin. Italics indicate that the feature cannot be observed. Scale bars: 1 µm. (D-F) enlargement of (A-C) green box. The inner tongue of the oligodendrocyte (white arrow) is only visible in the image recorded using interleaved scanning. Outside the axon, mitochondria (m) and other cellular membranes are observed. scale bars: 500 nm.

.
